# Neuropsychiatric sequelae in an experimental model of post-COVID syndrome in mice

**DOI:** 10.1101/2024.01.10.575003

**Authors:** Jordane Clarisse Pimenta, Vinícius Amorim Beltrami, Bruna Da Silva Oliveira, Celso Martins Queiroz-Junior, Jéssica Barsalini, Danielle Cunha Teixeira, Luiz Pedro de Souza-Costa, Anna Luiza Diniz Lima, Caroline Amaral Machado, Bárbara Zuccolotto Schneider Guimarães Parreira, Felipe Rocha da Silva Santos, Pedro Augusto Carvalho Costa, Larisse De Souza Barbosa Lacerda, Matheus Rodrigues Gonçalves, Ian de Meira Chaves, Manoela Gonzaga Gontijo Do Couto, Victor Rodrigues de Melo Costa, Natália Ribeiro Cabacinha Nóbrega, Bárbara Luísa Silva, Talita Fonseca, Filipe Resende, Natália Teixeira Wnuk, Hanna L. Umezu, Gabriel Campolina-Silva, Ana Cláudia dos Santos Pereira Andrade, Renato Santana de Aguiar, Guilherme Mattos Jardim Costa, Pedro Pires Goulart Guimarães, Glauber Santos Ferreira da Silva, Luciene Bruno Vieira, Vanessa Pinho, Antônio Lúcio Teixeira, Mauro Martins Teixeira, Aline Silva De Miranda, Vivian Vasconcelos Costa

## Abstract

The global impact of the COVID-19 pandemic has been unprecedented, and presently, the world is facing a new challenge known as Post-COVID syndrome (PCS). Current estimates suggest that more than 65 million people are grappling with PCS, encompassing several manifestations, including pulmonary, musculoskeletal, metabolic, and neuropsychiatric sequelae (cognitive and behavioral). The mechanisms underlying PCS remain unclear. The present study aimed to: (i) comprehensively characterize the acute effects of pulmonary inoculation of the betacoronavirus MHV-A59 in immunocompetent mice at clinical, cellular, and molecular levels; (ii) examine potential acute and long-term pulmonary, musculoskeletal, and neuropsychiatric sequelae induced by the betacoronavirus MHV-A59; and to (iii) assess sex-specific differences. Male and female C57Bl/6 mice were initially inoculated with varying viral titers (3×10³ to 3×10^5^ PFU/30 μL) of the betacoronavirus MHV-A59 via the intranasal route to define the highest inoculum capable of inducing disease without causing mortality. Further experiments were conducted with the 3×10^4^ PFU inoculum. Mice exhibited an altered neutrophil/lymphocyte ratio in the blood in the 2^nd^ and 5^th^ day post-infection (dpi). Marked lung lesions were characterized by hyperplasia of the alveolar walls, infiltration of polymorphonuclear leukocytes (PMN) and mononuclear leukocytes, hemorrhage, increased concentrations of CCL2, CCL3, CCL5, and CXCL1 chemokines, as well as high viral titers until the 5^th^ dpi. While these lung inflammatory signs resolved, other manifestations were observed up to the 60 dpi, including mild brain lesions with gliosis and hyperemic blood vessels, neuromuscular dysfunctions, anhedonic-like behavior, deficits in spatial working memory, and short-term aversive memory. These musculoskeletal and neuropsychiatric complications were exclusive to female mice and were prevented after ovariectomy. In summary, our study describes for the first time a novel sex-dependent model of PCS focused on neuropsychiatric and musculoskeletal disorders. This model provides a unique platform for future investigations regarding the effects of acute therapeutic interventions on the long-term sequelae unleashed by betacoronavirus infection.

## 1. Introduction

The global impact of the COVID-19 pandemic caused by the betacoronavirus SARS-CoV-2 has been unprecedented. It has affected over 699 millions of people, resulting in more than 6,9 millions of deaths worldwide (WHO, 2023). The peak of the pandemic overwhelmed the healthcare systems globally (Huang et al., 2020; Tan et al., 2020). Despite the subsequent development and approval of vaccines, our understanding of the enduring effects of COVID-19, particularly the long-term consequences known as long-COVID or Post-COVID syndrome (PCS), remains incomplete (Phillips and Williams, 2021).

Post-COVID syndrome (Proal et al., 2023; WHO, 2023) is a condition marked by the persistence of symptoms for months or even years after confirmed acute infection by SARS-CoV-2 (Proal and VanElzakker, 2021; WHO, 2023). Current estimates suggest that more than 65 million people are grappling with PCS (Ballering et al., 2022). The condition seems to affect approximately 10-30% of non-hospitalized individuals, 50-70% hospitalized individuals, and around 10-12% of vaccinated individuals (Davis et al., 2023). Post-COVID syndrome may affect any age group and is not necessarily associated with the severity of the acute phase of the disease (Davis et al., 2023; Long et al., 2020; Mao et al., 2020a; Mao et al.,2020b; Wiersinga et al., 2020; Zheng et al., 2020).

Clinically, PCS encompasses a broad spectrum of manifestations attributed to the widespread viral tropism associated with the expression of the ACE-2 receptor across various cell types (Hikmet et al., 2020). Symptoms of PCS include fatigue, impaired breathing (Tsuchida et al., 2023), arthralgia and myalgia (Romero et al., 2023), bone pain (Davis et al., 2021), cardiac symptoms as palpitations (Jiang et al., 2021), gastrointestinal changes including altered bowel habits and bloating (Comelli et al., 2022), as well as neuropsychiatric impairment, such as cognitive complaints (often referred to as “brain fog”), anxiety and depression (reviewed in Badenoch et al., 2022). These neuropsychiatric sequelae constitute a major concern since they may be associated with an increased risk of developing long-term cognitive impairment and dementia (García-González et al., 2023). A systematic review and meta-analysis examining persistent neuropsychiatric symptoms following COVID-19 revealed that among nearly 19,000 patients, 27.4% reported sleep disorder, 24.4% experienced fatigue, and 15.7% exhibited symptoms consistent with post-traumatic stress disorder (PTSD) (Simani et al., 2021), 35,5% cognitive dysfunction, 19.1% anxiety, 12.9% depression, 11.4% dysosmia, 7.4% dysgeusia, 6.6% headache, 5.5% disorder sensorimotor and 2.9% dizziness (Badenoch et al., 2022). Another comprehensive meta-analysis of 151 studies involving 1,285,407 participants across 32 countries examined the long-term physical and mental effects of COVID-19. Among the 659,454 survivors studied up to 12 months post-infection, 28.7% reported fatigue, 18.3% experienced depression, 17.9% showed symptoms of PTSD, and 19.7% displayed cognitive deficits (Zeng et al., 2023). Importantly, a study has shown that the risk of neuropsychiatric symptoms may be higher in COVID-19 than in other conditions, such as sepsis (Stallmach et al., 2022). Several hypotheses have been proposed to explain the development of neuropsychiatric and other non-pulmonary symptoms during the PCS, but the underlying mechanisms remain elusive. These hypotheses encompass: (i) the systemic acute inflammatory process and dysregulation of the immune response (Frere et al., 2022); (ii) the persistence of SARS-CoV-2 in immune privileged reservoirs (Stein et al., 2022); (iii) autoimmunity triggered by infection (Woodruff et al., 2022); (iv) microbiome dysbiosis (Mendes de Almeida et al., 2023); (v) reactivation of latent viruses such as Epstein-Barr (EBV) (Chen et al., 2021); (vi) dysfunctional neuronal signaling (Spudich and Nath, 2022); and/or (vii) sex-related susceptibility (Bai et al., 2022). Given the societal impact of the disease, there is a great need to understand the pathogenesis of PCS, which would allow the development of specific therapeutic strategies. Currently, the available approaches have focused on symptom management and rehabilitation measures (Möller et al., 2023).

Wild-type mice have shown resistance to SARS-CoV-2 infection (Dinnon et al., 2020; Gu et al., 2020). To study coronavirus infections in mice, researchers commonly use betacoronaviruses such as Murine Hepatitis Viruses (MHV-1, MHV-3, MHV-A59, and MHV-S strains), which naturally infect mice. They are associated with pulmonary infection and disease, effectively mimicking several aspects of human coronavirus infections (Andrade et al., 2021a; De Albuquerque et al., 2006; Yang et al., 2014). In this study, we thoroughly examined the acute effects of the intranasal inoculation of the betacoronavirus MHV-A59 and the potential long-term pulmonary and neuropsychiatric sequelae. Our findings revealed that MHV-A59 intranasal inoculation leads to transient lung infection and female hormone-dependent brain inflammation, followed by long-term cognitive and behavioral changes mimicking several aspects of long-COVID syndrome. This model provides a unique platform for investigating the pathogenesis of long-COVID and the therapeutic impact of antiviral, anti-inflammatory, or neuroprotective strategies.

## 2. Material and Methods

### 2.1 Mice

Animal experimental procedures were approved by the Ethical Committee for Animal Experimentation of the Universidade Federal de Minas Gerais (UFMG) (process number 140/2023). Experiments were carried out with male and/or female wild-type 5 and 6-weeks-old C57BL/6 mice (Central Animal House of the UFMG), as described in figure legends. Mice were housed in individually ventilated cages placed in an animal care facility at 24°C[±[2°C on a 12-h light/12-h dark cycle, receiving *ad libitum* access to water and food.

### 2.2 Cells and virus

L929 cells were cultured under a controlled atmosphere (37°C and 5% CO_2_) in high-glucose Dulbecco’s modified Eagle’s medium (DMEM) supplemented with 7% fetal bovine serum (FBS), 100 μg/mL streptomycin and 100 U/mL penicillin. The MHV-A59 strain was purchased from ATCC (Manassas, Virginia, USA) and propagated in L929 cells.

### 2.3 MHV-A59 infection

Mice were intraperitoneally anesthetized with ketamine (80 mg/kg) and xylazine (15 mg/kg) and received an intranasal inoculation of 30 μl sterile saline solution containing or not (Mock controls) MHV-A59 at different concentrations (3[×[10^3^ to 3[×[10^5^ PFU/30 µL), as described in figure legends. Weight loss and survival were monitored daily for up to 14 days post-inoculation (dpi).

### 2.4 Sample collection

Mice were intraperitoneally anesthetized with ketamine (80 mg/kg) and xylazine (15 mg/kg), and blood samples were collected from the abdominal vena cava for cell count (neutrophil to lymphocyte ratio (NLR) analyses determined with a Celltac MEK-6500K hemocytometer (Nihon Kohden) and euthanized by cervical dislocation. The lungs were collected, and the left lobe was fixed in 4% formaldehyde-buffered solution for histological analyses. The right lung lobes, brain, liver, and spleen were snap-frozen in liquid nitrogen and stored at −80 °C for viral titration, ELISA, and PCR assays. In another set of experiments whole lung and brain samples were collected for flow cytometry assays.

### 2.5 Viral titration

To evaluate viral titers, serially diluted virus suspension of plasma samples, lung, liver, spleen, and brain tissue homogenates (1:9 tissue to DMEM) were inoculated onto a confluent monolayer of L929 cells grown in 24-well plates. After gentle shaking for 1 h, samples were removed and replaced with DMEM containing 0.8% carboxymethylcellulose, 2% FBS, and 1% penicillin-streptomycin-glutamine and kept for 3 days at 37 °C and 5% CO_2_. Then, cells were fixed with 10% neutral buffered formalin solution for 2h and stained with 0.1% crystal violet. Virus titers were determined as PFU/mL or PFU/mg of tissue.

### 2.6 Histopathology

To evaluate lung and brain inflammatory scores, formaldehyde-fixed and paraffin-embedded lung and brain tissues were sectioned into 5-μm-thickness slices and stained with hematoxylin and eosin (H&E) or Masson’s trichrome stain. The inflammatory score in mice lungs and brain was blinded-determined by a pathologist (C.M.Q.-J.) (Pimenta et al. 2023). Lung inflammatory score encompasses (1) airway (0 to 4 points), (2) vascular (0 to 4 points), (3) parenchyma damage (0 to 5 points), and (4) general neutrophil infiltration (0 to 5 points).

In the brain, the evaluation was carried out in the cortex and hippocampus following a point scale: 0, no lesion; 1, mild tissue injury and/or mild inflammation; 2, mild tissue injury and/or moderate inflammation; 3, definitive tissue damage (neuronal loss and parenchymal damage) and intense inflammation; 4, necrosis (complete loss of all tissue elements with the presence of cellular debris). Meningeal inflammation was assessed using a point scale (0 to 4), with 0 representing no inflammation and 1 to 4 corresponding to 1 to 4 layers affected by inflammation. The final score was the sum of the cerebral cortex and hippocampus scores plus the meningitis score, totaling 12 points (Pimenta et al, 2023).

### 2.7 Mechanics of the respiratory system

To measure the compliance of the respiratory system, full-range pressure-volume (PV) curves were completed like those described in previous studies of our group (Pereira et al., 2023; Andrade et al., 2021) and adapted from Limjunyawong et al., 2015 and Robichaud et al., 2017. Briefly, mice were divided into three groups: control (mock), 5-and 30-dpi. The animals were deeply anesthetized, and a polyethylene tube (P50) was inserted into the mouse’s trachea. The PV curve was generated by injecting air volume continuously using a 3 mL syringe and an automated syringe pump (Bonther, Ribeirão Preto, SP, Brazil) at a rate of 3 mL/min until the intratracheal pressure peaked at approximately 35 cm H_2_O. At peak pressure, the syringe pump was manually switched for the deflation limb, deflated at the same rate until the pressure reached approximately −15 cm H_2_O, and finally inflated again to the resting lung volume. Both volume and pressure signals were acquired and recorded using PowerLab software (LabChart v7, AdInstruments, Sydney, Australia). Inflation and deflation were repeated at least twice to obtain accurate curves, and full-range PV curves for each animal were obtained. If leaks or high pressures were detected, the data for that animal were not included in the analysis. Vital capacity was determined by maximum insufflation (lung volume at 35 cm H_2_O), and static compliance of the respiratory system (expressed in mL/cmH2O) was measured at the steepest point of the deflation limb of the PV curve (Andrade et al., 2021; Limjunyawong et al., 2015; Robichaud et al., 2017).

### 2.8 Cytokine and chemokine measurement

For the determination of inflammatory mediators in the lung and brain tissues, the samples were homogenized in chilled cytokine extraction buffer (100 mM Tris [pH 7.4], 150 MM NaCl, 1 mM EGTA, 1 mM EDTA, 1% Triton X-100, 0.5% sodium deoxycholate and 1% protease inhibitor cocktail). After centrifugation (10,000 rpm, 10 min, 4 °C), the supernatant was collected and used to measure inflammatory mediators using the DuoSet enzyme-limited immunosorbent assay (ELISA) system (R&D Systems). The concentrations of CXCL1, CCL2, CCL3, CCL5, TNF, IL-1β, IL-6, IFN-γ, IL-10 and TGF-β were measured in the lung supernatant. IL-6, IFN-γ, BDNF (brain-derived neurotrophic factor), and CX3CL1 (fractalkine) were measured in the prefrontal cortex (PFC) and hippocampus.

To perform plasma estradiol measurement, 0.3 mL of blood was collected and centrifuged at 2,000 rpm for 10 min to separate the plasma, which was subsequently stored at −20°C. Plasma estradiol levels were measured using enzyme-linked immunosorbent assay (ELISA) with the commercial AccuBind kit (Monobind Inc., USA). To determine the estradiol concentration, a curve graph was constructed using the absorbance of each duplicate serum reference and the corresponding concentration of the estradiol standard in pg/mL.

The quantification of anti-MHV-A59 IgG/IgM antibodies followed the methodology outlined by Costa et al. (2014). In brief, 96-well plates were initially coated with the MHV-A59 isolate at 10[ PFU/well. Subsequently, the plates underwent a one-hour exposure to UV-C light to render the virus inactive, followed by an overnight incubation at 4°C and 1% bovine albumin PBS solution blocking for a duration of 2 hours. Then, serum samples diluted at 1:200 for IgG and 1:20 for IgM were applied and allowed to incubate for 3 hours at 37 °C. Post-incubation, a peroxidase-Anti-IgG or anti-IgM antibody was applied and incubated for an additional 2 hours. Results were quantified in abstract units, derived from the detected absorbance readings in conjunction with the respective dilutions.

Measurements of the absorbance were conducted using an ELISA reader (Status-labsystems, Multiskan RC, Uniscience do Brasil) set at 490 nm. The antibodies used are listed in Table 1.

**Table 1.**
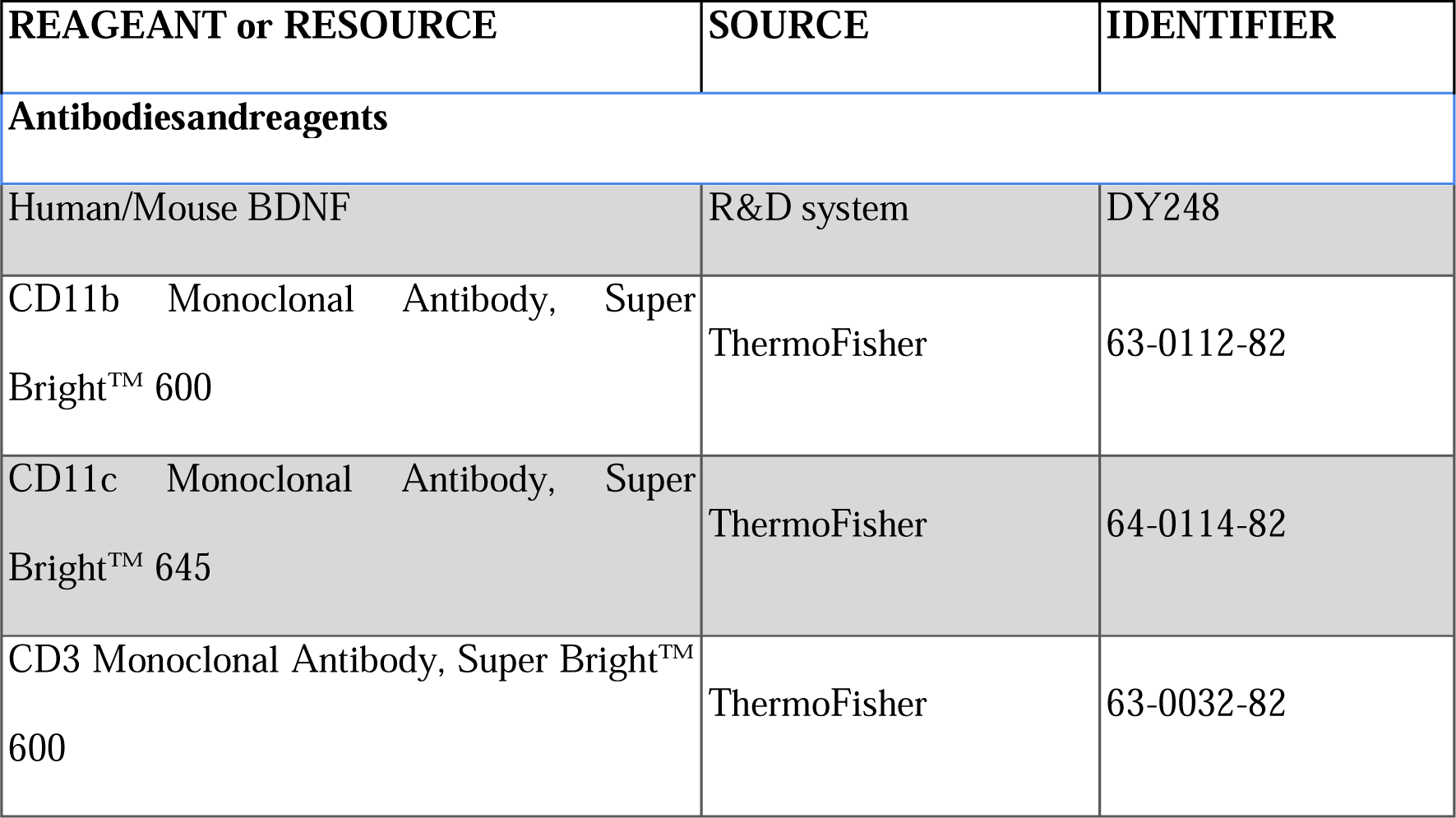
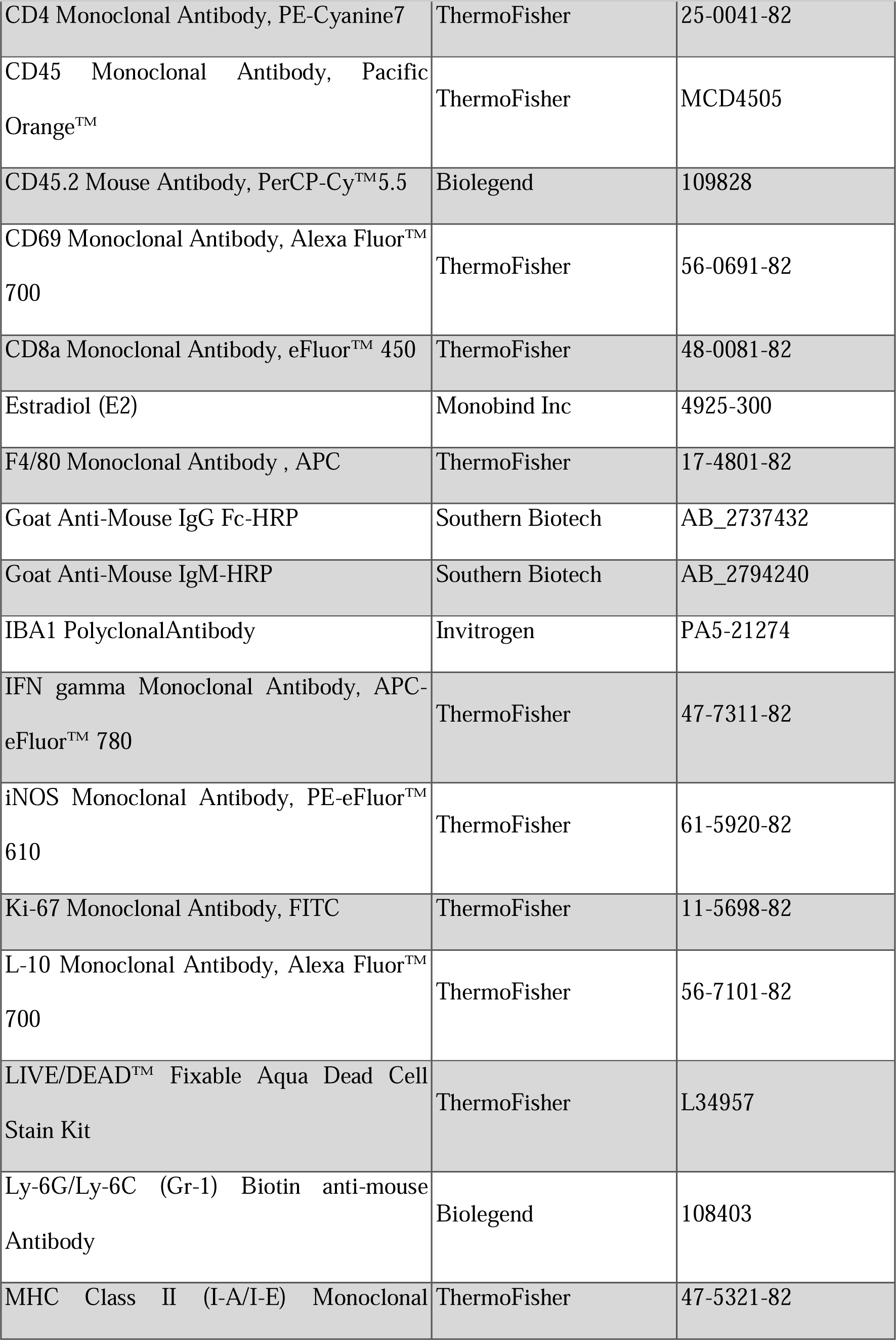
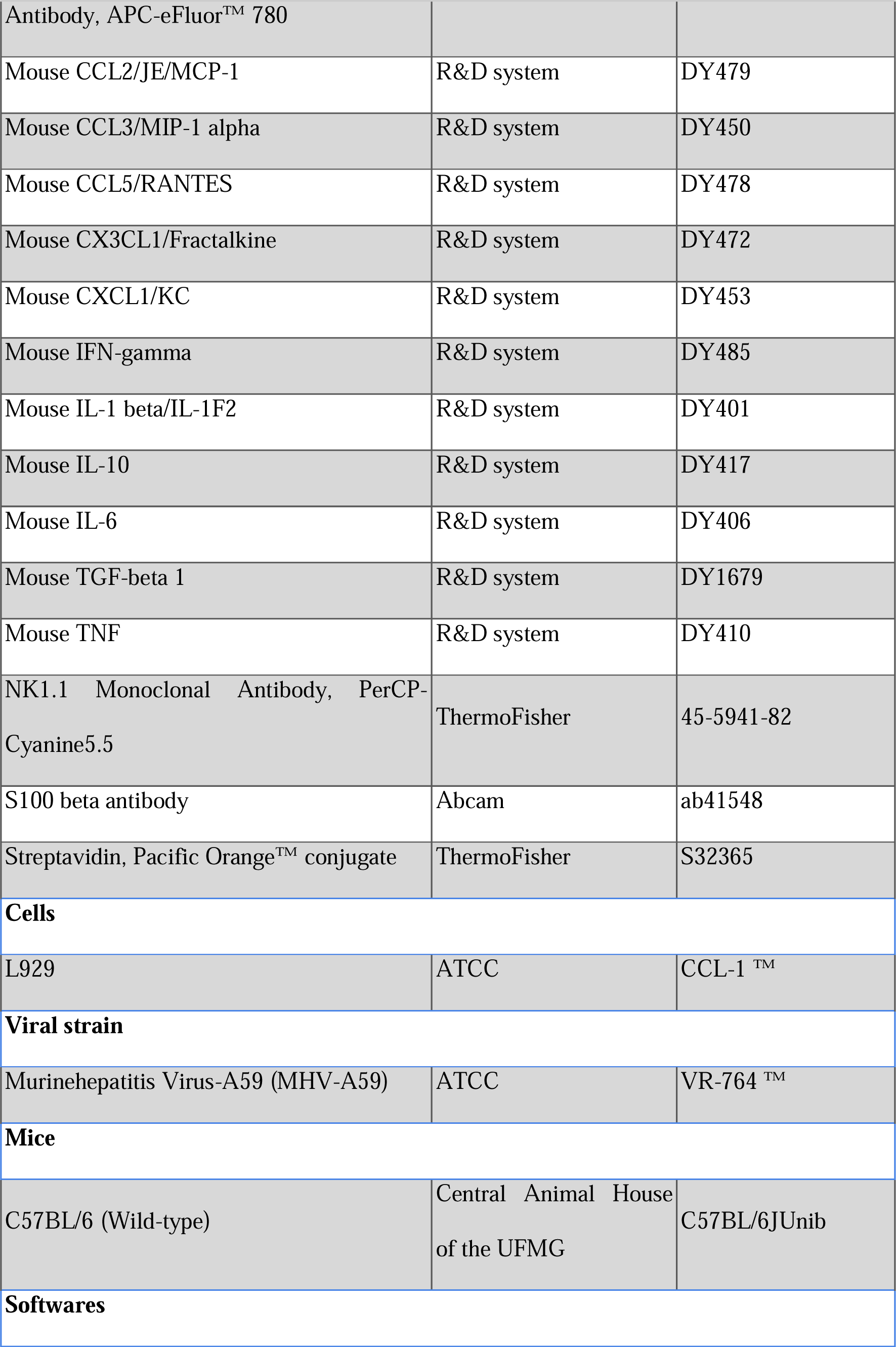
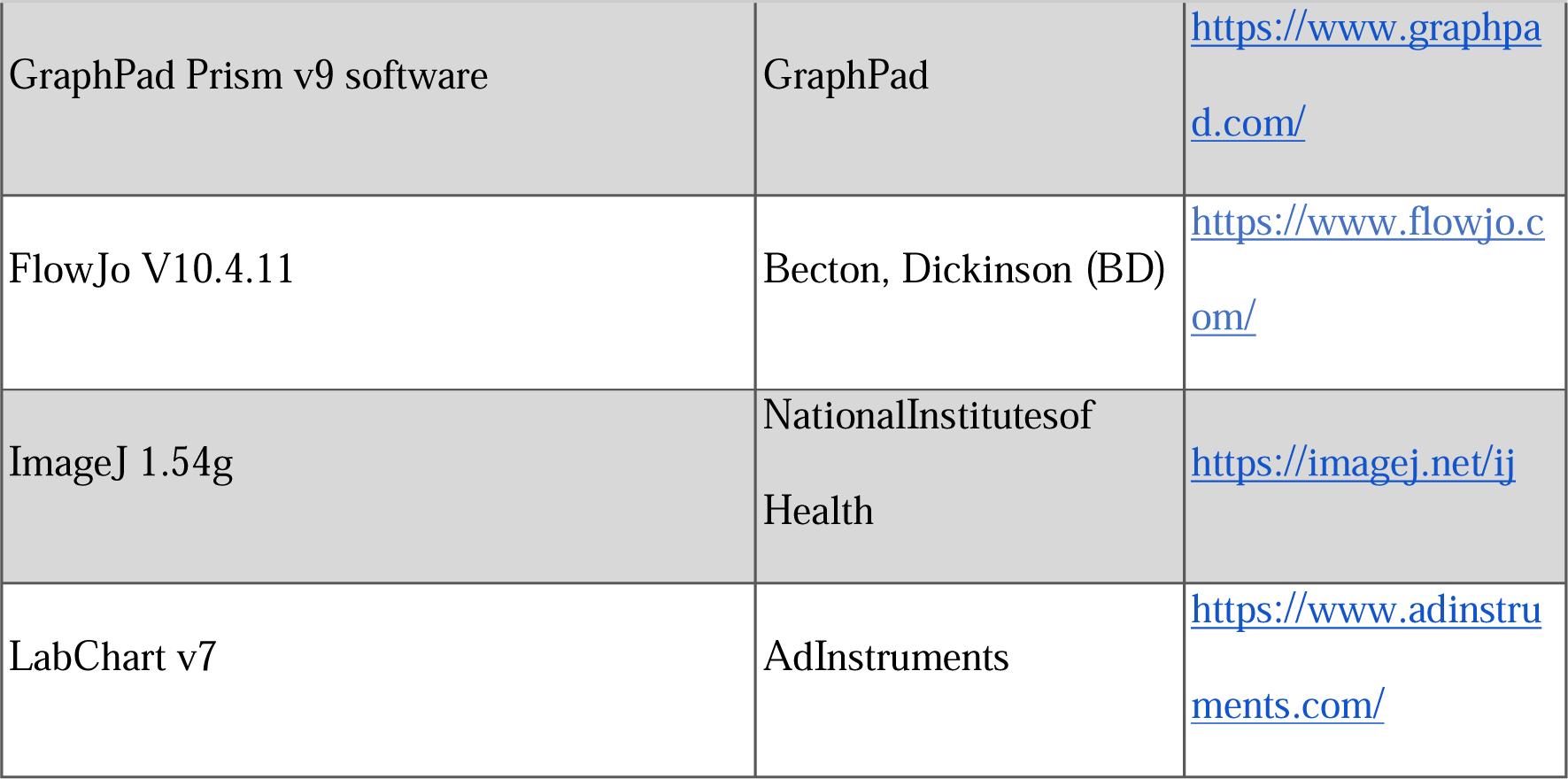
Antibodies, reagents, cells, viral strain, mice and softwares.

### 2.9 Flow cytometry

To assess infiltrating cell immunophenotyping and intracellular cytokines, brain and lung tissues were collected, processed, and enriched. Brain tissues underwent maceration, Percoll gradient separation, and filtration. Lung tissues were dissected, digested with Collagenase I, dissociated, and filtered. Cells were washed in the FACS buffer, and dead cells were excluded. Extracellular and intracellular antigens were labeled after fixation and permeabilization. The LSR-FORTESSA equipment was used for acquisition, and data were analyzed using singlets with FSC-A versus FSC-H gate. In the lung, live leukocytes (neutrophils, T CD4^+^, CD8^+^, NK cells, and dendritic cells) were characterized. In the brain, infiltrated T subsets and neutrophils were evaluated, along with activated microglia. IFN-γ, IL-10, and iNOS production were measured in each cell subset in the brain and lung microenvironment using intracellular staining. Isolated cells were maintained for 4 hours in RPMI with supplements, and Brefeldin A. FlowJo V10.4.11 was used for data analysis, and the antibodies used are listed in Table 1.

### 2.10 IBA-1 e S100B Immunohistochemistry in brain samples

Sections of the cerebral cortex and hippocampus from MHV-A59-infected mice and the control group (mock) were quantitatively analyzed for microglia, IBA-1 (ionized calcium-binding adapter molecule 1; antibody PA5-21274, Invitrogen; 1:150) and astrocytes (anti-S100 beta antibody; ab41548, Abcam; 1:75), according to the manufacturer’s instructions (Vector Elite kit). The quantification of IBA-1 and S100B was carried out from 10 random sections of the histological section of the brain areas using the Image J® software (National Institute of Health, USA).

### 2.11 Purification of synaptosomes

Mice infected or not with MHV-A59 were euthanized by cervical dislocation. The cortex and/or hippocampus were promptly dissected and homogenized in a gradient solution composed of 320 mM sucrose, 0.25 mM dithiothreitol, and 1 mM EDTA. Subsequently, the homogenate underwent low-speed centrifugation (1000 *g* × 10 min). Synaptosomes were then isolated from the supernatant through discontinuous Percoll density gradient centrifugation, as detailed by Dunkley et al. (1988). The isolated nerve terminals were resuspended in Krebs–Ringer-HEPES (KRH) solution, which included 124 mM NaCl, 4 mM KCl, 1.2 mM MgSO_4_, 10 mM glucose, 25 mM HEPES, with pH adjusted to 7.4 and devoid of additional CaCl_2_, achieving a concentration of approximately 10 mg/mL. For the assessment of glutamate release and intrasynaptosomal calcium concentration, 30 μL aliquots were prepared and maintained on ice until further use.

### 2.12 Glutamate measurements

To assess continuous glutamate release, a fluorometric assay was conducted using a Synergy TM2 fluorimeter (Biotek®). Fluorescence emission was recorded with an excitation wavelength of 360 nm and emission at 450 nm. Glutamate release was indirectly quantified by monitoring the fluorescence increase resulting from NADPH production in the presence of glutamate dehydrogenase type II and NADP^+^ (according to Nicholls et al., 1987). In brief, synaptosomes were incubated with 1 mM CaCl_2_ and 1 mM NADP^+^ in KRH medium for 5 minutes. Glutamate dehydrogenase (50 units per well) was introduced after 5 minutes. Depolarization was induced using 33 mM KCl. A calibration curve was established by adding glutamate (5 nM/μL) to the reaction medium. Glutamate levels were normalized to the total protein content per well.

### 2.13 Intrasynaptosomal free calcium measurements

For the determination of intrasynaptosomal free calcium concentration, synaptosomes were preincubated with 5 μmol/L of Fura-2 pentakis (acetoxymethyl) Ester (FURA2-AM) probe for 30 minutes at 35.5 °C. Subsequently, the synaptosomes were centrifuged (3,000 *g* × 60 s), resuspended in KRH without CaCl_2_, and reincubated for 30 minutes. Following washout with CaCl_2_-free KRH, synaptosomes were promptly employed for the quantification of intracellular free calcium ([Ca^2+^]i). Fluorescence was recorded at an excitation wavelength of 340/380 nm and an emission of 510 nm. CaCl_2_ (1 mmol/L, final concentration) was introduced into the synaptosomal suspension before reading, and 33 mM of KCl was added to induce calcium influx. Finally, 10% SDS was added to establish Rmax, and tris-EGTA (3 mol/L Tris, 400 mmol/L EGTA, pH 8.6) was introduced to establish Rmin, following the procedures outlined by Grynkiewicz et al., 1985; Nicholls et al., 1987.

### 2.14 Ovariectomy (OVX)

To study the effect of female sex hormones in MHV-A59-induced sequelae, 6-week-old female C57BL/6 mice were anesthetized (ketamine 80 mg/kg, xylazine 15mg/kg, i.p.) and ovariectomized bilaterally (group OVX). Sham-operated mice were subjected to the same experimental procedure but without ovariectomy (group SHAM). Fourteen days after the surgery, groups of Sham or OVX mice were inoculated with MHV-A59 (3 × 10^4^ PFU in 30 μL) or sterile saline (30 μL) intranasally. Mice were evaluated for neuropsychiatric sequelae along 60 dpi or euthanized at 2, 5, or 60 dpi for lung, brain, uterus, and plasma collection. The uterus weight was determined to detect uterus atrophy induced by female hormones’ deficiency.

### 2.15 Open field test

The open field test was employed 5, 16, and 28 days after MHV-A59 infection to measure spontaneous locomotor activity as described elsewhere (Hefner and Holmes, 2007). Briefly, mice were gently placed in an arena (30 cm × 30 cm × 50 cm square), and then they were allowed to explore the arena for 30 min freely. Key parameters were recorded, such as spontaneous locomotor activity and the percentage of time spent in the center of the arena (a measure of anxiety-like behavior). The test was conducted using the Phenotyper apparatus (Noldus, Information Technology, Leesburg, VA, USA), containing a digital video camera and infrared lights in each arena. The EthoVision video tracking software (EthoVision XT, Noldus Information Technology, Leesburg, VA, USA) was used for data recording and analysis.

### 2.16 Marble burying

The marble burying test was performed 34 days after MHV-A59 infection to evaluate compulsive-like behavior as described by Kalueff et al., 2006. Briefly, mice were placed in a rectangular cage (30 × 30 × 50 cm) with 20 cm of fresh bedding and 25 marbles placed equidistant to each other. The animals were allowed to explore and bury the marbles freely for 30 minutes. At the end of the session, we removed the mouse and measured the number of marbles buried. We considered only the balls buried by more than 2.5 cm. A significant increase in the number of balls buried indicates a compulsive-like behavior. However, associated with tests like the sucrose preference, a decrease in the number of balls buried reinforces an anhedonic-like behavior.

### 2.17 Sucrose preference test

The sucrose preference test (SPT) was performed to measure anhedonia-like behavior as described elsewhere (De Bundel et al., 2013). Anhedonia is conceived as the reduction or loss of interest in pleasurable feelings or enjoyable activities. The SPT is based on a two-bottle choice test that measures the preference of mice to intake a sweet solution, being the reduced interest indicative of anhedonia-like behavior. Double-housing mice were initially habituated for 2 days with two water bottles. During the next 3 days, one water bottle was replaced with a bottle with sucrose solution (1%). The percentage of sucrose consummation was recorded, and a significant decrease in this parameter indicates an anhedonic-like behavior.

### 2.18 Y maze test

The Y maze test was conducted 60 days after the MHV-59 infection to evaluate spatial working memory as described by Ribeiro et al., 2023. The Y-maze is composed of three arms intersecting at an angle of 120°. Mice were gently placed in the apparatus and allowed to explore the maze freely for 5 minutes. Following each trial, the maze was meticulously cleaned with a 70% alcohol solution and dried using paper towels. The percentage of spontaneous alternations between arms for 5 minutes was obtained through the index: [alternate total/ (total of entries in arms −2) × 100]. A significant decrease in spontaneous alternation indicates an impairment in spatial working memory.

### 2.19 Grip-force test

Grip-force in the forelimbs and all limbs was measured using Grip Strength Meter (Insight Equipamentos, Ribeirão Preto, SP) as previously described (Rossi et al., 2023). Mice were allowed to grasp the smooth pull bar with forelimbs and all limbs by holding their tails. Mice were then pulled backward in the horizontal plane, and the peak force (*g*) applied to the bar was recorded. Three trials were performed per mouse within the same session, and the average of the three trials was recorded. The average of these three trials was then converted into force using the formula F = m × g, where “F” represents force, “m” is the mass, and “*g*” is the acceleration due to gravity.

### 2.20 Step-down inhibitory avoidance test

The step-down inhibitory avoidance test was performed to assess short and long-term aversive memory 60 days after MHV-A59 infection, as previously described by Rodrigues et al., 2011. Briefly, in the training trial, mice were placed on the platform, and their latency to step down on the grid with all four paws was measured. Immediately after stepping down on the grid with the four paws, the animals received a single mild foot shock (0.2 mA, 2.0 s). A retention test trial was performed at 1.5 h (short-term aversive memory) and 24 h (long-term aversive memory) after the training section. In retention tests, each animal was placed on the platform again, and no shock was applied when the animal stepped down on the grid. The results were expressed as a latency period to step down the platform, with a cutoff of 180 seconds.

### 2.21 Olfactory discrimination test

Olfactory memory was assessed by the olfactory discrimination test at 6 and 38 days after MHV-A59 infection, as previously described by Prediger et al., 2010. The apparatus used for this test was an acrylic box divided into two equal compartments by an open door (7.0 cm²) in the center, enabling the animal to select between two compartments. One compartment, known as the familiar compartment, contained sawdust that had remained unchanged for a minimum of 3 days. Conversely, the other compartment, considered non-familiar, was filled with fresh sawdust. The animals were allowed to explore the environment freely for five minutes. The time spent in each compartment was recorded and subsequently analyzed using the EthoVision XT software (Noldus, Technology, Leesburg, VA, USA). The olfactory discriminative memory impairment was indicated by a significant decrease in the time spent in the familiar compartment.

### 2.22 PCR

RNA extraction from tissues was carried out using QIAamp®C Viral RNA kits, in accordance with the manufacturer’s instructions with specific adaptations. Tissue homogenization was performed in conjunction with the lysis buffer. The quantification of the extracted RNA was conducted using a spectrophotometer (NanoDrop™C, Thermo Scientific). For cDNA synthesis, the iScript™C gDNA Clear cDNA Synthesis Kit (BIO-RAD) was employed. The initiation of cDNA synthesis utilized 500 ng of total RNA, following the manufacturer’s protocols. The resulting total cDNA was previously diluted at a ratio of 1:10 for subsequent use in the qPCR assay. Fast SYBR™C Green Master Mix (Applied Biosystems™C) was employed along with primers at a concentration of 5nM. The forward primer sequence was 5′-CAGATCCTTGATGATGGCGTAGT-3′, and the reverse primer sequence was 5′-AGAGTGTCCTATCCCGACTTTCTC-3′. Additionally, RNA extraction was performed on a known PFU quantity of MHV-A59 to establish the standard, and the results were expressed in Arbitrary Units.

### 2.23 Statistical analyses

GraphPad Prism software (v.9.3.0) was used for the statistical analyzes. Significant outliers identified by the ROUT test (Q = 1%) were excluded from the data prior to subsequent analyzes. The data were tested for normality using the Shapiro-Wilk test. For data that passed the normality test, a confidence interval of95% was assumed. Two-way ANOVA and Dunnet’s multiple comparison test were used to analyze differences in weight changes and differences between mock controls and infected mice of respective sex at different time points. Simple Kaplan-Meier survival analysis was used to analyze the differences in survival probability. Student’s t-test (for normal distribution) or Mann–Whitney test (for non-normal distribution) was used to analyze the differences between the mock controls and the infected mice at a single time point. Two-way ANOVA and the Šídák multiple comparison test were used to analyze the differences between the sexes at the respective infection time points. For data that did not pass the normality test, a confidence interval of 99% was assumed for Dunnet’s multiple comparison test or the Šídák multiple comparison. Three-way ANOVA and Tukey’s multiple comparison tests were used to analyze the differences between sham or ovariectomized (OVX) mock or infected mice at different time points. For data that did not pass the normality test, a confidence interval of 99% was assumed for the Tukey multiple comparison test. Data are shown as mean values ± standard error of the mean.

## 3. Results

### 3.1 Intranasal instillation of MHV-A59 induces a transient pulmonary infection and an inflammatory state

Intranasal MHV-A59 infection induces acute pneumonia and severe lung injuries in C57BL/6 wild-type mice (Yang et al., 2014). However, comprehensive analyses of the chronic and systemic consequences associated with MHV-A59 infection have not been conducted. Here, we intranasally inoculated 5-week-old C57BL/6 mice with MHV-A59 at different viral loads (3 × 10^3^, 3 × 10^4^, or 3 × 10^5^ PFU/30 μL) (Fig. 1A). Infection with MHV-A59 reduced the body weight of mice in an inoculum-dependent manner (Fig. 1B), whereas only the 3 × 10^5^PFU inoculum was lethal to infected mice (Fig. 1C). To investigate the systemic, pulmonary, and cerebral changes induced by MHV-A59 infection, we opted for a higher, non-lethal inoculum of 3 × 10^4^ PFU to proceed with the experiments. Mice were euthanized at intervals of 2, 5, 8-, 16-, 30-, and 60-days post-infection (dpi), and samples of lung, liver, spleen, and blood were subsequently collected (Fig. 1A).

**Figure 1.**
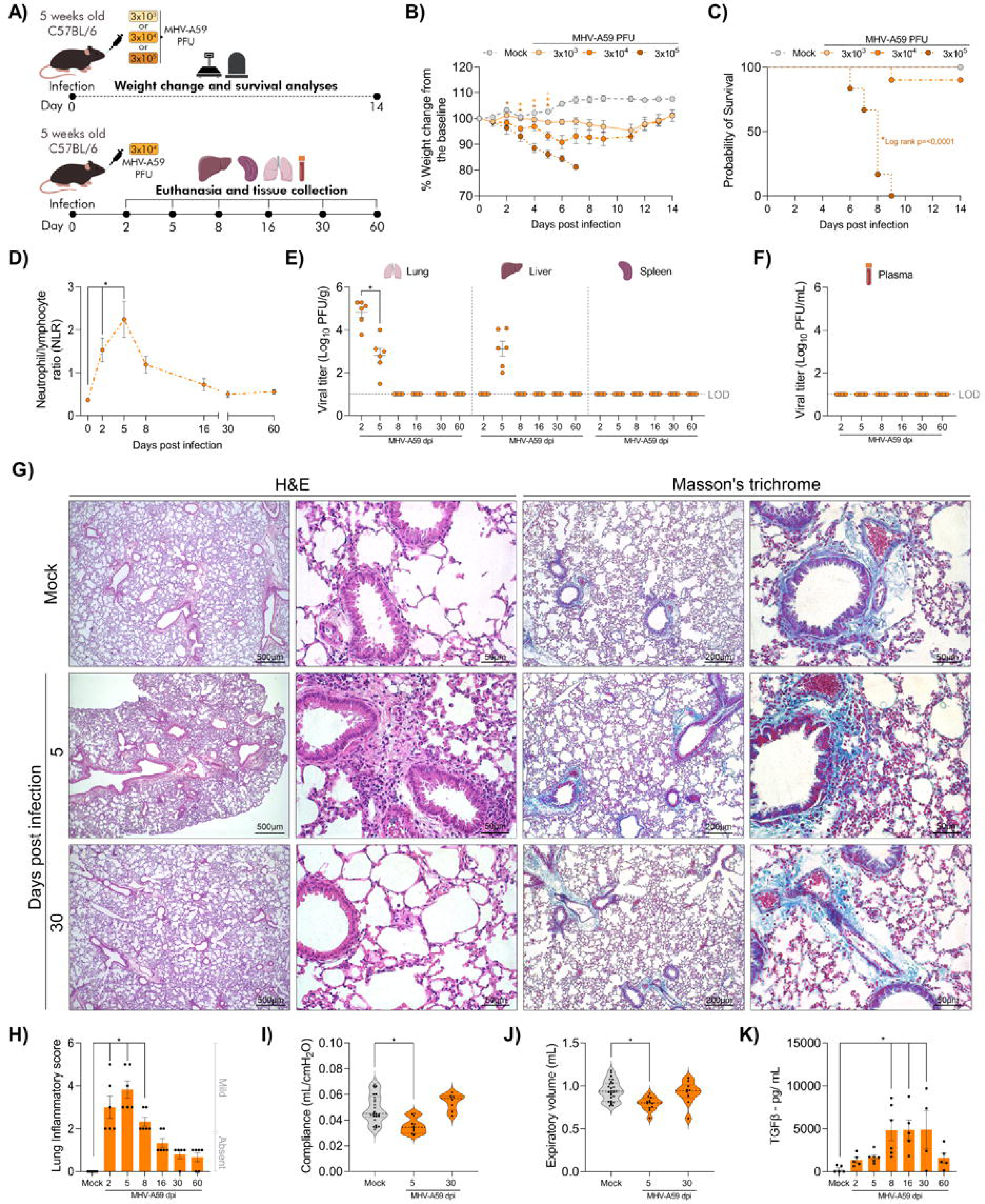
Intranasal instillation of MHV-A59 in mice results in transitory lung injury and function impairment. (A) Experimental design. (B) Body weight percentage changes upon infection with 3×10^3^, 3×10^4^ or 3×10^5^ PFU/30 μL of MHV-A59 versus mock controls. Significance was determined by Two-way ANOVA and Dunnet’s multiple comparison test *p < 0.05 (n = 8-10). (C) Survival curve of MHV-A59-infected mice versus mock controls. Significance was determined by Simple Kaplan-Meier survival analysis *p < 0.05 (n = 6-10). (D) Blood neutrophil for lymphocyte ratio (NLR). Significance was determined by Two-way ANOVA and Dunnet’s multiple comparison test *p < 0.01 (n = 6). Viral load determined in lung, liver, and spleen extracts (E), and plasma (F) of MHV-A59-infected mice by plaque assay. The results are presented as log_10_ PFU per gram of tissue or milliliter of plasma. Significance was determined by Student’s t-test *p < 0.05 (n = 6). (G) Hematoxylin and eosin (H&E) and Masson’s trichrome staining of lung sections of mock controls and 5 and 30-dpi MHV-A59-infected mice. (H) Histopathological assessment in relation to overall lung inflammatory score. Significance was determined by Two-way ANOVA and Dunnet’s multiple comparison test *p < 0.01 (n = 5-6). (I) Analysis of total lung volume (vital capacity) and (J) pulmonary compliance of mock controls and 5 and 30-dpi MHV-A59 infected mice. Significance was determined by Two-way ANOVA and Dunnet’s multiple comparison test *p < 0.05 (n = 9-28). (K) Lung tissue levels of TGF-β. Significance was determined by Two-way ANOVA and Dunnet’s multiple comparison test *p < 0.05 (n = 4-6). L.O.D. (limit of detection).

Coronavirus infection can lead to an imbalance in the neutrophil-to-lymphocyte ratio (NLR), a potential indicator of the severity and mortality of the disease (Henry et al., 2020; Liao et al., 2020; Ponti et al., 2020). Mice infected with MHV-A59 exhibited increased NLR in the blood at 2 and 5-dpi (Fig. 1D). As MHV-A59 and SARS-CoV–2 have been shown to spread to extrapulmonary sites (Körner et al., 2020; Synowiec et al., 2021), our investigation assessed viral titers not only in lung samples but also in plasma, liver, and spleen obtained from MHV-A59-infected mice. The intranasal inoculation of MHV-A59 resulted in a transient lung infection, with the peak viral titer at 2 dpi. The titer decreased by 5 dpi and remained undetectable after that (Fig. 1E). Replicating virus was also detected in the liver (Fig. 1E), with mild histopathological alterations (Figure S1 A-B).

As shown by the H&E-stained slides, intranasal infection with MHV-A59 resulted in transient mild lung inflammation characterized by perivascular and peribronchiolar leukocyte infiltrate, desquamation of bronchiolar cells, hyperplasia of the alveolar walls and also points of hemorrhage in some samples (Fig. 1H). In addition, infected mice presented impaired lung function, with a significant reduction in both vital capacity (Fig. 1I) and compliance of the respiratory system (Fig. 1J) at 5dpi. This stiff behavior of the respiratory system is consistent with pulmonary restrictive disease (Mortola 2019) and can be associated with fibrosis (Schaller et al., 2020). As TGF-β is a pivotal regulator of collagen deposition (Meng et al., 2016), we investigated the concentration of this cytokine in the lungs of MHV-A59-infected mice. Infection with MHV-A59 increased the TGF-β levels in the lung at 8, 16, and 30 dpi (Fig. 1K). Despite the elevated TGF-β levels, there was no accumulation of collagen in the lungs of the mice throughout the infection, as evidenced by Masson’s trichrome-stained slides (Fig. 1G).

### 3.2 MHV-A59 intranasal infection promotes differential accumulation of pulmonary chemokines and leukocytes

Next, we evaluated the molecular and cellular profiles involved in MHV-A59-induced lung inflammation (Fig. 2). Considering that biological sex has an impact on both immune response and COVID-19 outcomes (Scully et al., 2020; Takahashi et al., 2020), we stratified the sexes in our analyses to elucidate potential mechanisms contributing to the disease phenotype. Both male and female MHV-A59-infected mice showed elevated levels of CCL2, CCL3, and CCL5 in lung tissue, while only male-infected mice showed elevated levels of CXCL1 (Fig. 2A). Additionally, we assessed the levels of the pro-inflammatory cytokines TNF, IFN-γ, IL-6, and the anti-inflammatory cytokine IL-10 in the lung tissue of MHV-A59-infected mice. However, these levels were not statistically different between non-infected and infected mice (data not shown). At 5 dpi, both female and male MHV-A59-infected mice displayed increased leukocyte accumulation in lung tissue (Fig. 2B and S2). While the levels of CXCL-1 were elevated in the lung tissue of male MHV-A59-infected mice, there was no alteration in neutrophil accumulation throughout the course of the disease in both female and male mice (Fig. 2C).

**Figure 2.**
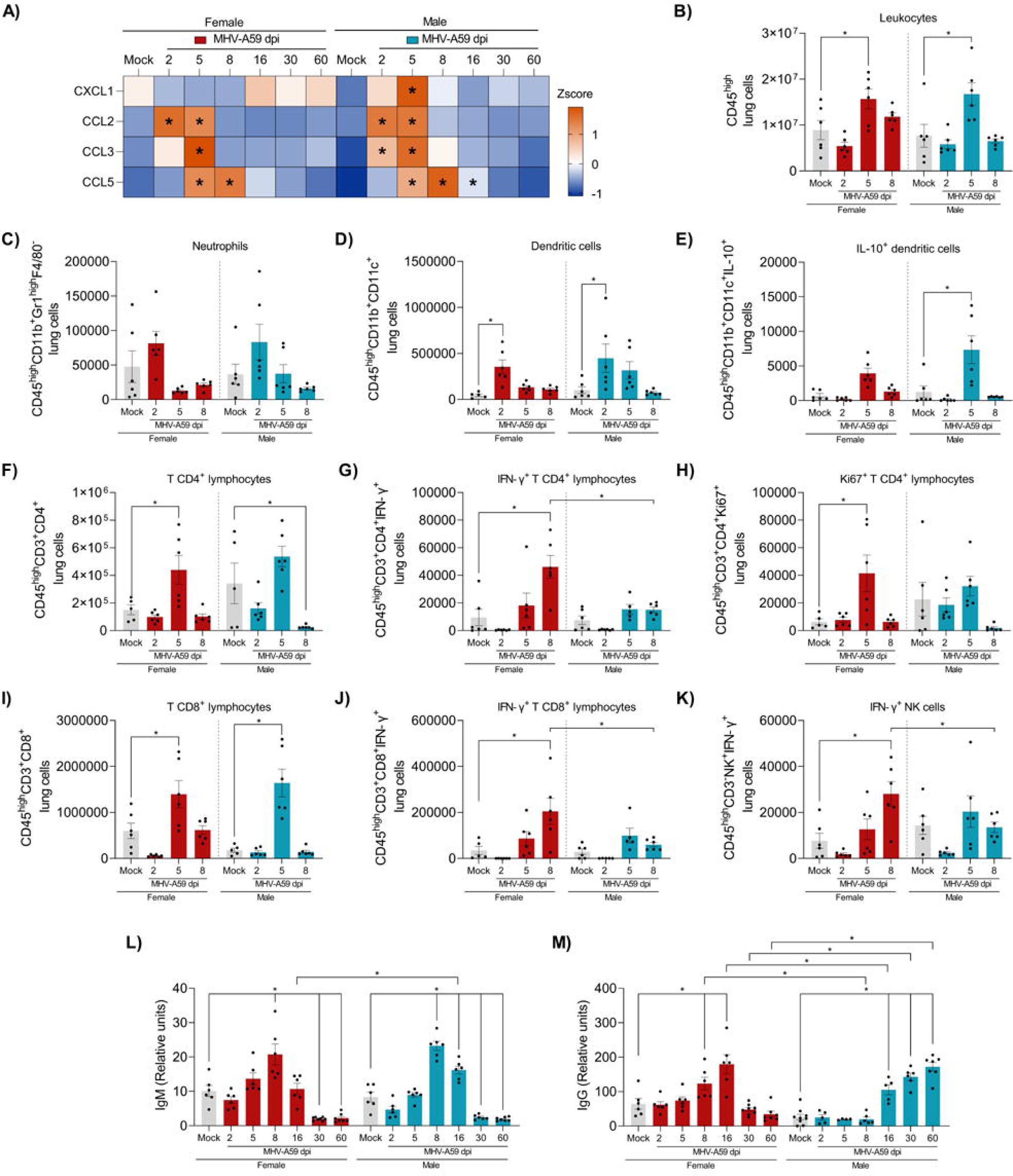
Intranasal infection with MHV-A59 promotes differential accumulation of pulmonary chemokines and leukocytes. **(A)** Z Score representation of pulmonary levels of CXCL1, CCL2, CCL3, CCL5 measured by ELISA assay. Significance was determined by Two-way ANOVA and Dunnet’s multiple comparison test *p < 0.01 (n = 4-6). Number of pulmonary leukocytes **(B)**, neutrophils **(C)**, dendritic cells **(D)**, IL-10^+^ dendritic cells **(E)**, CD4^+^ T lymphocytes **(F)**, Ki67^+^ CD4^+^ T lymphocytes **(G)**, IFN-γ^+^ CD4^+^ T lymphocytes **(H)**, CD8^+^ T lymphocytes **(I)**, IFN-γ^+^ CD8^+^ T lymphocytes **(J)**, and IFN-γ^+^ NK cells **(K)**, assessed by flow cytometry. Significance was determined by Two-way ANOVA and Dunnet’s multiple comparison test to analyze the differences between mock controls and infected mice of respective sex at different time points. Significance was determined by Two-way ANOVA and the Šídák multiple comparison test to analyze the differences between the sexes at the respective infection time points. *p < 0.05 to data that passed Shapiro-Wilk test and *p < 0.01 to data that did not pass Shapiro-Wilk test (n = 5-6). Plasma levels of IgM **(L)** and IgG **(M)**. Significance was determined by Two-way ANOVA and Dunnet’s multiple comparison test to analyze the differences between mock controls and infected mice of respective sex at different time points. Significance was determined by Two-way ANOVA and the Šídák multiple comparison test to analyze the differences between the sexes at the respective infection time points *p < 0.05 (n = 6-8).

Subsequently, at 5 dpi, a distinct subset of anti-inflammatory dendritic cells, marked by elevated IL-10 levels, aggregated in the lung tissue of MHV-A59-infected mice. Notably, this particular population was more abundant in male mice compared to their female counterparts (Fig. 2D-E). MHV-A59 induced a lymphocytic response in lung tissue, particularly notable in female animals. At 5 dpi, female infected mice showed an increased number of T CD4^+^ lymphocytes in the lungs (Fig. 2F). At 8 dpi, a pro-inflammatory subset of these cells, characterized by elevated levels of IFN-γ, accumulated in female infected mice but not in males (Fig. 2G). The T CD4-^+^ lymphocytes were characterized by high levels of the cell proliferation marker Ki67^+^ (Fig. 2H). Both female and male MHV-A59-infected mice exhibited an accumulation of T CD8^+^ lymphocytes in lung tissue (Fig. 2I). However, only female infected mice demonstrated an increase in the IFN-γ^+^ secreting CD8^+^ T lymphocyte subset in the lungs (Fig. 2J). While the overall number of pulmonary NK cells remained consistent between female and male mice throughout the disease (data not shown), a subpopulation of NK cells with high IFN-γ levels increased solely in the lung tissue of female infected mice (Fig. 2K) Protective humoral immunity against SARS-CoV-2 is a crucial determinant in COVID-19, correlating with clinical outcomes (Carrillo et al., 2021). Thus, we subsequently examined IgM and IgG levels against MHV-A59 in the plasma of infected mice. Plasma IgM and IgG levels were elevated in both female and male infected mice. Notably, males exhibited prolonged IgM plasma levels compared to females, whereas females demonstrated faster IgG production compared to male infected mice (Fig. 2L-M).

In sum, both female and male infected mice displayed the production of inflammatory mediators and the accumulation of cells in lung tissue following MHV-A59 infection. However, males exhibited increased chemokine production, while females demonstrated higher T lymphocyte activation.

### 3.3 MHV-A59-infected mice display behavioral and cognitive alterations in a time and sex-dependent manner

Following the characterization of lung effects induced by MHV-A59, we explored potential neuropsychiatric sequelae associated with Post-COVID syndrome (Xu et al., 2022). We analyzed behavioral and cognitive changes in male and female mice infected with MHV-A59 (Fig. 3A). In the open field test, no significant difference was found at 5 dpi (peak of lung disease) regardless of sex. Meanwhile, female MHV-A59 mice exhibited a significant decrease in spontaneous locomotor activity at 16 dpi. Notably, female mice exhibited complete recovery at 28 dpi, as evidenced by the total distance traveled in the open field. MHV-A59 infection did not result in locomotor activity impairment in male mice (Fig. 3B-D).

**Figure 3.**
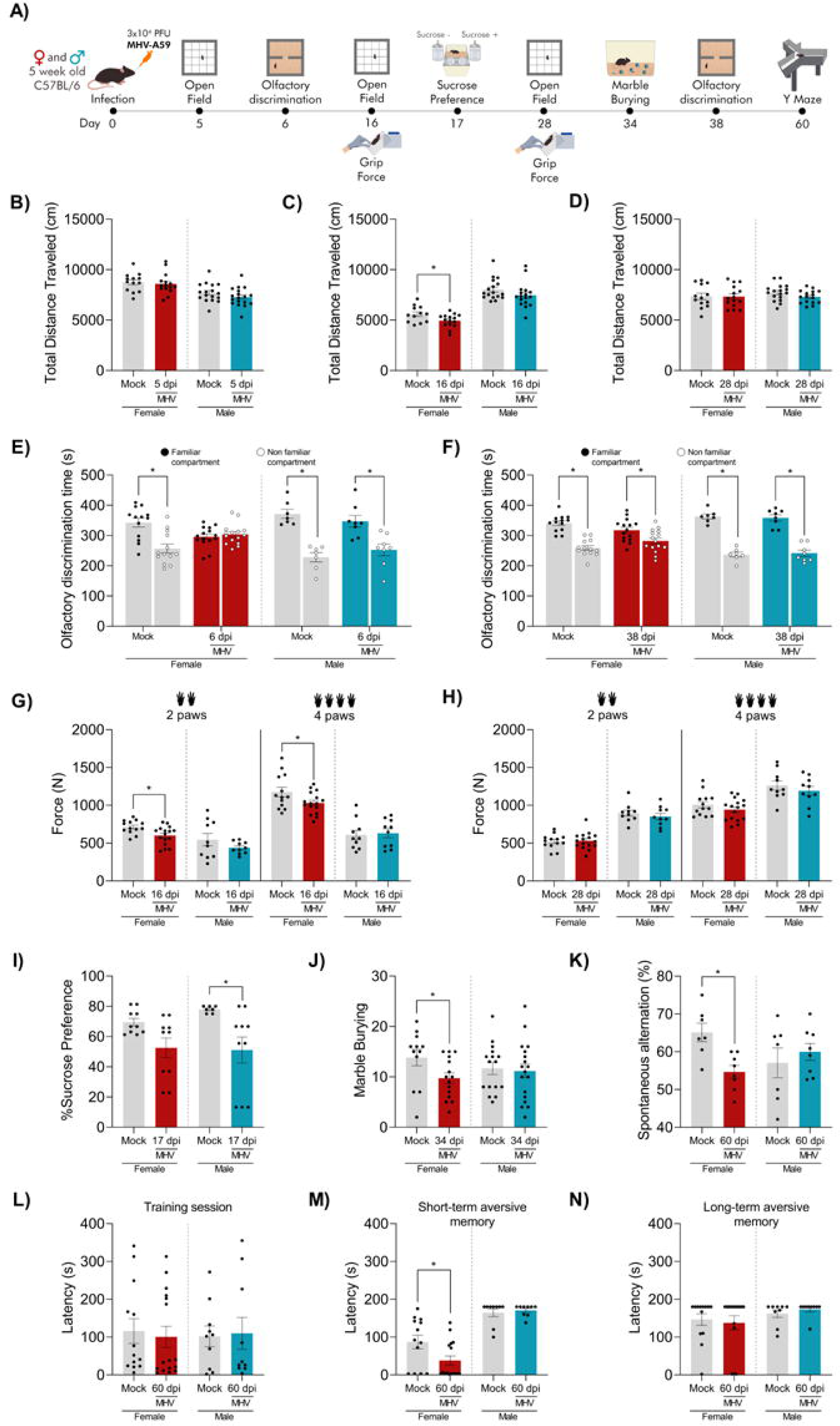
MHV-A59 induces behavioral and cognitive alterations in a time and sex dependent manner. **(A)** Experimental design. Total distance traveled in the open field test at **(B)** 5-, **(C)**16-, and **(D)** 28-dpi (n = 13-18). Olfactory discrimination time **(E)** 6- and **(F)** 38-dpi (n = 7-15). Force in newtons (N) in 2 or 4 paws grip force test at **(G)** 16- and **(H)** 28-dpi (n = 10-16). **(I)** Sucrose preference test at 17-dpi (n = 10-12). **(J)** Marble burying test at 34-dpi(n = 13-18). **(K)** Spontaneous alternation in Y-maze test at 60-dpi (n = 7-8). **(L)** Latency in training session, **(M)** short-term aversive memory, and **(N)** long-term aversive memory, in Y-maze test at 60-dpi (n = 10-16). Significance was determined by Student’s t-test to data that passed Shapiro-Wilk test and Mann–Whitney test to data that did not pass Shapiro-Wilk test *p < 0.05.

As olfactory loss is a core symptom of COVID-19 (Harapan and Yoo, 2021), we investigated olfactory discrimination memory at early and later time points after MHV-A59 infection. Male-infected mice had no disturbance in the olfactory discrimination memory, while at 6 dpi female-infected mice presented a significant dysfunction compared to mock groups (Fig. 3E). Olfactory discrimination memory dysfunction was completely resolved by 38 dpi (Fig. 3F). Neuromuscular dysfunction is also an important symptom related to COVID-19 (Rossi et al., 2023). At 16 dpi, both female MHV-A59-infected mice displayed a significant decrease in the forelimb and for all limbs grip force compared with mock controls (Fig. 3G). Grip strength impairments were not observed for forelimbs and all limbs at 28 dpi regardless of sex (Fig. 3H).

Male MHV-A59-infected mice displayed an anhedonic-like behavior, as indicated by a significant decrease in the percentage of sucrose preference at 17 dpi (Fig. 3I). A reduction in the percentage of sucrose preference was also observed in infected female mice (Fig. 3I), however it did not reach statistical significance (p= 0.056). Interestingly, female animals presented a decrease in the number of buried marbles at 34 dpi, also suggesting an anhedonic-like behavior (Fig. 3J).

Importantly, only female mice showed significant cognitive dysfunctions at 60 dpi MHV-A59 infection. There was a significant decrease in the percentage of spontaneous alternations in the Y maze, indicating an impairment in spatial working memory. No significant differences were found between MHV-A59 infected male mice and mock controls (Fig. 3K). Regarding aversive memory, mock and infected mice displayed similar step-down latency in the training session regardless of sex (Fig. 3L). MHV-A59 female mice showed impairment in short-term but not long-term aversive memory compared with mocks, as indicated by a decrease in the step-down latency 1.5 h but not 24 h after the training session. No significant changes in aversive memory were observed in male mice after MHV-A59 infection (Fig. 3L-N). Therefore, female mice presented more significant signs of chronic neuropsychiatric sequelae after MHV-A59 infection.

### 3.4 MHV-A59 infection induces prominent neurochemical and cellular alterations in the brain of female mice

Given the behavioral and cognitive changes identified post MHV-A59 infection, particularly in female mice, we proceeded to investigate the biochemical and morphological alterations induced by the infection in the brain(Fig. 4A). MHV-A59 RNA viral copies were found in the brains of infected animals at from 2-to 8-dpi, in both sexes. However, the replicative virus was not detected by plaque assay (data not shown). The highest numbers of virus RNA copies were detected on days 5 and 8 dpi, with females having notably more virus at 5 dpi compared to males (Fig. 4B).

**Figure 4.**
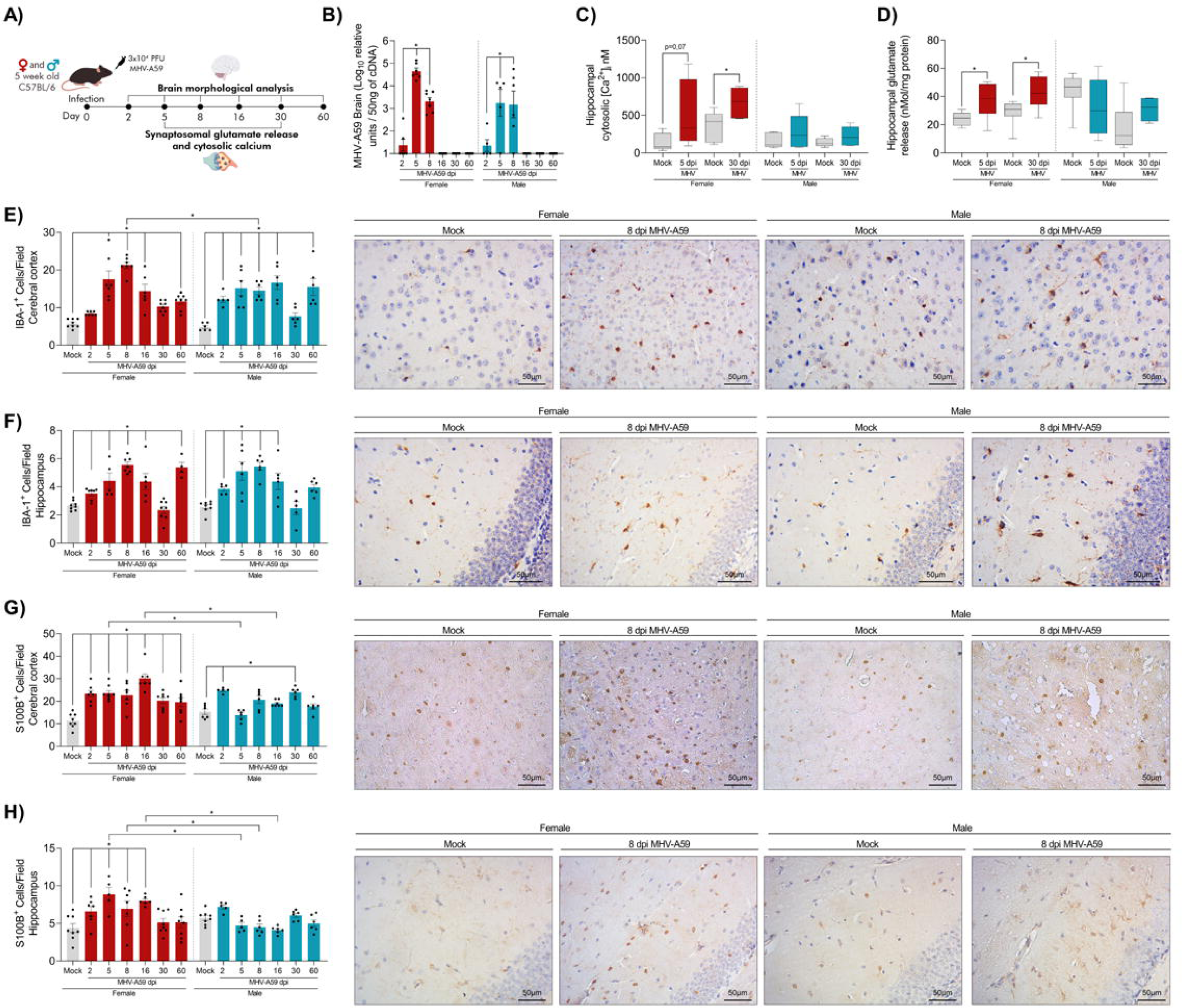
MHV-A59 infection induces more prominent neurochemical and cellular alterations in females. **(A)** Experimental design. **(B)** MHV-A59 brain viral load measured by PCR. Significance was determined by Two-way ANOVA and Dunnet’s multiple comparison test to analyze the differences between infected mice of each sex at different time points or by Two-way ANOVA and the Šídák multiple comparison test to analyze the differences between the sexes at the respective infection time points *p < 0.01 (n = 5-7). **(C)** Hippocampalintrasynaptosomalcalciumconcentration (n = 5-7) and**(D)** glutamate release(n = 6-8). Significance was determined by Student’s t-test to data that passed Shapiro-Wilk test and Mann–Whitney test to data that did not pass Shapiro-Wilk test *p < 0.05. **(E)** IBA-1^+^ cells in the cerebral cortex (n = 6-8) and **(F)** hippocampus (n = 4-8). **(G)** S100B^+^ cells in the cerebral cortex (n =5-8) and **(H)** hippocampus (n = 5-8). Significance was determined by Two-way ANOVA and Dunnet’s multiple comparison test to analyze the differences between mock controls and infected mice of respective sex at different time points or by Two-way ANOVA and the Šídák multiple comparison test to analyze the differences between the sexes at the respective infection time points. *p < 0.05 to data that passed Shapiro-Wilk test and *p < 0.01 to data that did not pass Shapiro-Wilk test.

A potential mechanism associated with brain damage is neuronal excitotoxicity (Verma et al., 2022). To examine this hypothesis, we assessed intracellular calcium and glutamate levels in isolated nerve terminals, specifically synaptosomes isolated from the hippocampus (Fig. 4C and D, respectively) and cortex (Fig S3A-B, respectively) of both female and male mice at 5- and 30-dpi. Female, but not male mice, exhibited higher levels in intrasynaptosomal calcium concentrations (Fig. 4C) and an increase in hippocampal glutamate levels (Fig. 4D). As for the cortex, no changes were found in glutamate and calcium levels in both sexes (Fig S3A-B). These findings underscore the sex-specific profile in the neurochemical response to MHV-59 infection.

Histopathological analysis of the cerebral cortex revealed discrete changes in both sexes, such as the presence of leukocytes surrounding some hyperemic vessels (Fig S3C-D). Immunohistochemical assays showed an increase in the number of IBA-1^+^ (microglia/macrophages) cells in the brain cortex after MHV-A59 infection in both sexes (Fig. 4E). In males, this increase was detected from the 2^nd^ to the 60^th^-dpi when compared to the mock group, while females showed an increase from the 5^th^-dpi onwards (Fig 4E). Noteworthy, at 8-dpi, females showed higher numbers of IBA1^+^ cells when compared to males (Fig. 4E). IBA-1 labeling was also performed in the hippocampus and there was a similar profile of IBA-1^+^ cells in both sexes. Relative to their respective mocks, there was an increase in cells labeled for IBA-1 from 2-to 60-dpi (Fig. 4F). The number of S100B^+^ astrocytes increased in the brain cortex of female mice after the 2^nd^-dpi onwards when compared to the mock group. In males, this increase was only observed on the 2^nd^-dpi (Fig. 4G). Significant difference was observed between females and males at 5 and 16-dpi, with females showing higher numbers than males (Fig. 4G). A similar pattern was detected in the hippocampus (Fig. 4H). Females showed a significant increase in S100B^+^ cells from 2-to 16-dpi when compared to their control group, while no increase was observed in males. When comparing the sexes, females exhibited higher numbers of hippocampus S100B^+^ cells at 5-, 8-, and 16-dpi than males (Fig. 4H). The figures demonstrate IBA-1^+^ and S100B^+^ cells in the cerebral cortex and hippocampus at the peak of MHV-A59 infection compared to their respective controls.

### 3.5 MHV-A59 intranasal infection triggers differential accumulation of brain leukocytes and inflammatory mediators

Next, we evaluated the cellular and molecular profiles involved in MHV-A59-induced CNS inflammation (Fig S4). Both female and male mice infected with MHV-A59 exhibited increased leukocyte accumulation in the brain at 8-dpi when compared to respective controls (Fig. 5A). The number of neutrophils increased in the brains of females at 8-dpi, but not in males (Fig. 5B). Additionally, IBA-1^+^ cells, indicative of microglia, revealed a pattern of active cells (CD45^int^CD11b^+^F4/80^+^MHCII^+^) in both sexes also at 8-dpi when compared to the mock groups (Fig. 5C). The expression of iNOS in this population of activated microglia was upregulated in females and was significantly higher when compared to the male group (Fig. 5D).

**Figure 5.**
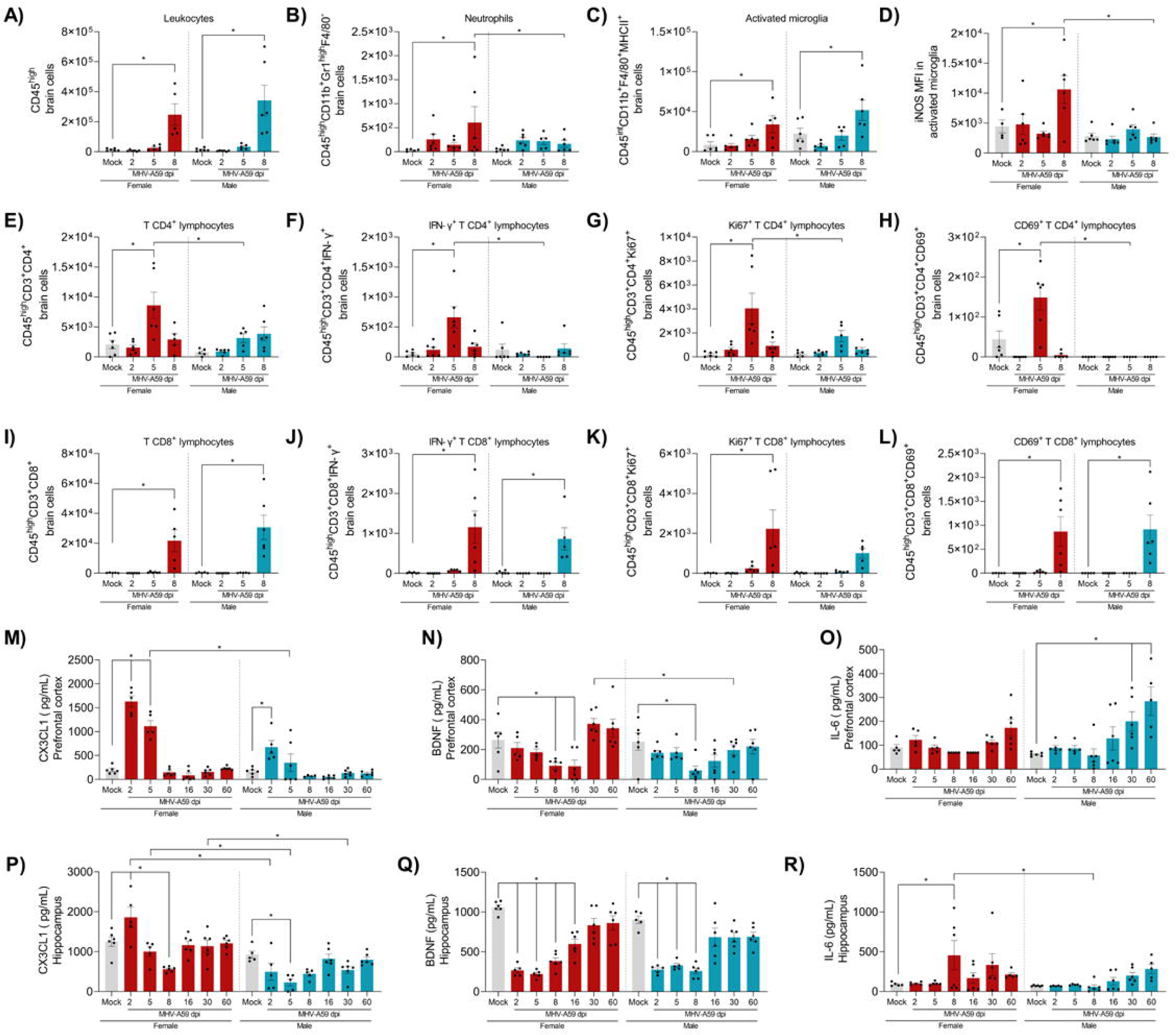
MHV-A59 infection triggers differential accumulation of leukocytes and inflammatory mediators in the brain. **(A)** Number of brain leukocytes, **(B)** neutrophils, **(C)** activated microglia, **(D)** iNOS MFI in activated microglia, **(E)** CD4^+^ T lymphocytes, **(F)** INF-γ^+^ CD4^+^ T lymphocytes, **(G)** Ki67^+^ CD4^+^ T lymphocytes, **(H)** CD69^+^ CD4^+^ T lymphocytes, **(I)** CD8^+^ T lymphocytes, **(J)** IFNy^+^ CD8^+^ T lymphocytes, **(K)** Ki67^+^ CD8^+^ T lymphocytes, and **(L)** CD69^+^ CD8^+^ T lymphocytes, assessed by flow cytometry. Significance was determined by Two-way ANOVA and Dunnet’s multiple comparison test to analyze the differences between mock controls and infected mice of respective sex at different time points and by Two-way ANOVA and the Šídák multiple comparison test to analyze the differences between the sexes at the respective infection time points. *p < 0.05 to data that passed Shapiro-Wilk test and *p < 0.01 to data that did not pass Shapiro-Wilk test (n = 5-6). Prefrontal cortex levels of **(M)** CX3CL1, **(N)** BDNF, and **(O)** IL-6, and hippocampal levels of **(P)** CX3CL1, **(Q)** BDNF, and **(R)** IL-6, measured by ELISA assay. Significance was determined by Two-way ANOVA and Dunnet’s multiple comparison test to analyze the differences between mock controls and infected mice of respective sex at different time points and by Two-way ANOVA and the Šídákmultiple comparison test to analyze the differences between the sexes at the respective infection time points. *p < 0.05 to data that passed Shapiro-Wilk test and *p < 0.01 to data that did not pass Shapiro-Wilk test (n = 5-6).

MHV-A59 infection also prompted a lymphocytic response in brain tissue, which was prominent in infected female mice since they exhibited higher numbers of T CD4^+^ lymphocytes at 5-dpi than their control group and males at the same time point (Fig. 5E). Additionally, at 5-dpi, there was an accumulation of a pro-inflammatory subpopulation of T CD4^+^ lymphocytes in the brain of infected female mice, expressing CD69^+^, which is a marker of early activation, Ki67^+^, a marker of cell proliferation, aside from high levels of IFN-γ (Fig. 5F-H). All of these cell markers were significantly increased when compared to males.

Regarding the subpopulation of T CD8^+^ cells, there was an expansion of these cells in the brains of both male and female infected mice at 8-dpi (Fig. 5 I). Similarly, there was an increase in the number of IFN-γ-producing CD8^+^ T cells and T CD8^+^CD69^+^ cells, although the number of T CD8^+^ Ki67^+^ cells only increased in infected females when compared to the control group and male mice (Fig. 5J-L).

In line with the leukocyte infiltration, MHV-A59 induced an increase in CX3CL1 in the prefrontal cortex (PFC) of females at 2- and 5-dpi when compared to the mock group, while in males, such increase occurred only at 2-dpi. Furthermore, this increase was significantly more pronounced in females (Fig. 5M). Females also showed a reduction in BDNF levels at the 8^th^ and 16^th^-dpi in the PFC, while males showed this reduction only at 2 dpi. At the 30^th^ - dpi, there was a significant difference between the sexes, with males exhibiting lower levels of BDNF than females (Fig. 5N). In contrast, male mice infected with MHV-A59 showed an increase in IL-6 levels in the PFC at 30- and 60-dpi, while infected females showed no alterations (Fig. 5O).

MHV-A59 infection also led to an increase at 2-dpi and a reduction at 8-dpi in hippocampus levels of CX3CL1 in female mice. In contrast, CX3CL1 levels in the hippocampus of infected male mice decreased at 5-dpi when compared to mock and were significantly lower than females at 2-, 5-, and 30-dpi (Fig. 5P). Hippocampus BDNF levels decreased at 2-, 5-, 8-, and 16-dpi in infected females and at 2-, 5-, and 8-dpi in infected males, compared to their respective control groups. Nevertheless, no sex-based differences were observed (Fig. 5Q). Quantifying IL-6 levels in the hippocampus indicated a peak in infected female mice at 8-dpi, contrasting with the mock group. Furthermore, a significant increase in IL-6 was observed in infected females compared to males at this time point (Fig. 5R). Quantification of the cytokine IFN-γ was carried out in the same way as other mediators. However, it was not detected at the protein level.

### 3.6 MHV-A59-induced CNS dysfunction is dependent on female hormones

In line with the current results, women seem to be more clinically affected by Post-COVID syndrome (Phillips and Williams, 2021). To explore this sex-hormone dependent phenotype, we initially assessed estradiol serum levels in MHV-A59 infected female mice. Intranasal MHV-A59 infection did not induce changes in estradiol levels during the acute phase of the disease. However, these levels surged nearly sevenfold in the long-term compared to the age-matched control group (Fig. 6A). The next step was to study the disease and its sequelae in a female hormones-deficient context. Thus, we ovariectomized (OVX) mice and infected them with MHV-A59 to analyze the role of sex hormones in this infectious process (Fig. 6 B). Assessment of viral load in the lungs of SHAM-infected and OVX-infected animals revealed similar amounts of virus at 2 and 5 dpi (Fig. 6C).

**Figure 6.**
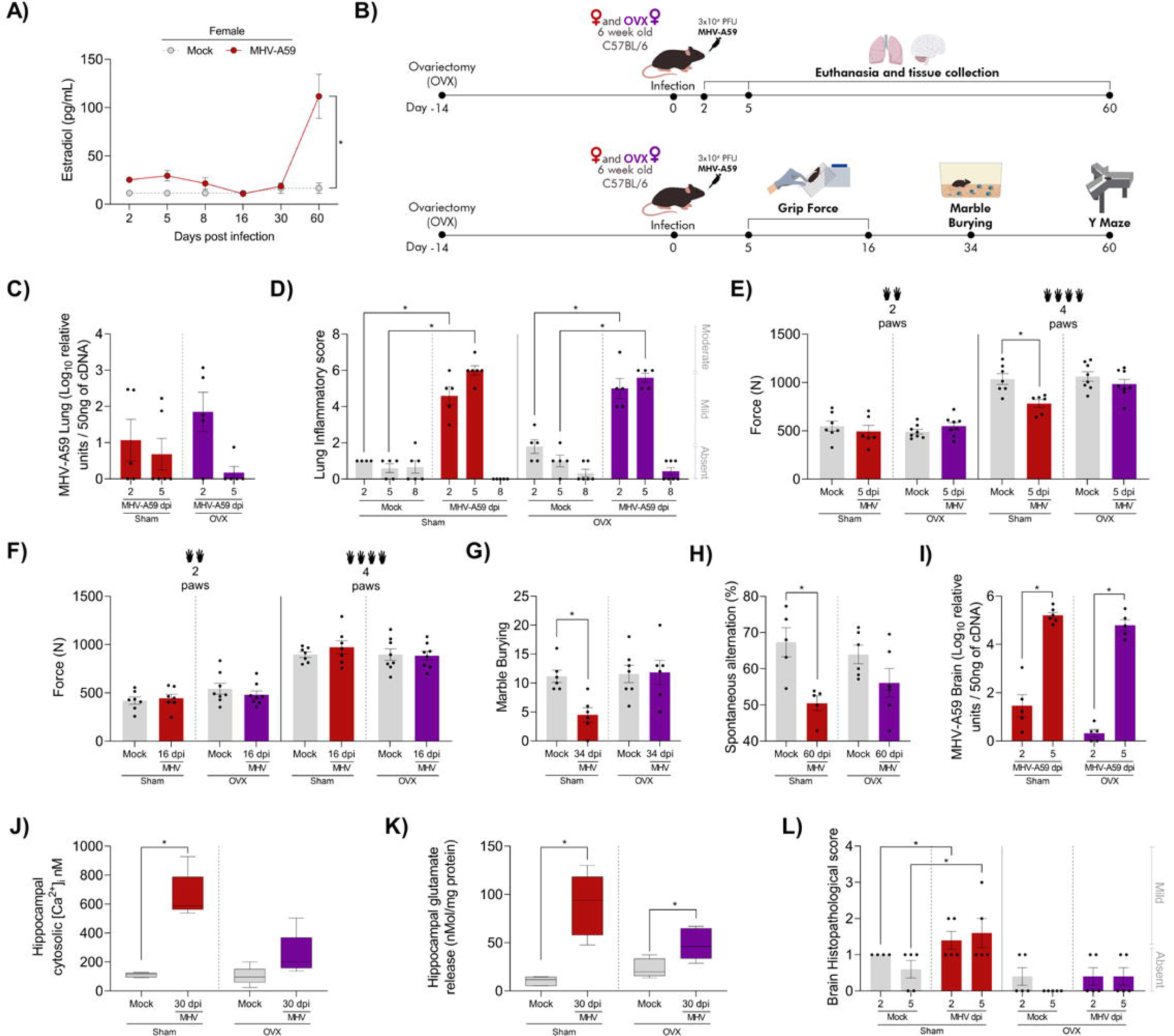
Ovariectomy (OVX) reduces histopathological brain damage, prevents behavioral alterations and minimizes neurochemical imbalance in the hippocampus induced by MHV-A59 infection. **(A)** Plasma estradiol levels in mock and MHV-A59-infected female mice. Significance was determined by Two-way ANOVA and the Šídák multiple comparison test to analyze the differences between mock controls and MHV-A59 infected female mice at the respective time points. *p < 0.05 (n = 3-11). **(B)** Experimental design. **(C)** MHV-A59 lung viral load measured by PCR. Significance was determined by Two-way ANOVA and Dunnet’s multiple comparison test *p < 0.01 (n = 5-6) **(D)** Overall lung inflammatory score. Significance was determined by Three-way ANOVA and Tukey’s multiple comparison test to analyze the differences between sham or OVX, mock or infected mice, at different time points *p < 0.01 (n = 4-7). Force in newtons (N) in 2 or 4 paws grip force test **(E)** 5- and **(F)** 16-days post-infection (n = 6-8). **(G)** Marble burying test 34 days post-infection (n = 6-7). **(H)** Spontaneous alternation in Y-maze test 60 days post-infection (n = 5-6). Significance was determined by Student’s t-test to data that passed Shapiro-Wilk test and Mann–Whitney test to data that did not pass Shapiro-Wilk test *p < 0.05. **(I)** MHV-A59 brain viral load measured by PCR. Significance was determined by Two-way ANOVA and Dunnet’s multiple comparison test *p < 0.01 (n = 5-6) **(J)** Hippocampal intrasynaptosomal calcium concentration (n = 5) and **(K)** glutamate release(n = 5-7). Significance was determined by Student’s t-test to data that passed Shapiro-Wilk test and Mann–Whitney test to data that did not pass Shapiro-Wilk test *p < 0.05. **(L)** Overall brain histopathological score (n = 4-5). Significance was determined by Three-way ANOVA and Tukey’s multiple comparison test to analyze the differences between sham or OVX, mock or infected mice, at different time points *p < 0.01.

The histopathological lung damage of infected females subjected to OVX was similar to that of SHAM mice, both at 2- and 5-dpi (Fig. 6D and S5). Nevertheless, female hormone deficiency in the OVX group protected mice when evaluating the neuropsychiatric sequelae induced by the infection. While MHV-A59 infection triggered a significant reduction in grip strength in all limbs at 5-dpi (Fig. 6E), OVX mice had no alterations in the grip strength of the limbs in the acute phase of infection (Fig. 6E). No significant changes in neuromuscular function were found 16 days after MHV-A59 infection (Figure 6F). Importantly, ovariectomy also prevented the MHV-A59-associated anhedonia-like behavior and spatial working memory deficit at 34- and 60-dpi, respectively (Fig. 6 G-I).

Regarding viral load in the brain at 2 e 5 dpi, SHAM and OVX animals did not show significant differences (Fig. 6I). Accordingly, calcium concentration in the hippocampus of SHAM-infected animals increased relative to their control group at 30 dpi, while OVX-infected mice exhibited no significant alterations at the same time point (Fig. 6J). In contrast, glutamate release increased similarly in the hippocampus of both SHAM and OVX-infected groups (Fig. 6K). Accordingly, OVX prevented the mild histopathological damage induced by MHV-A59 in brain tissue compared with the SHAM-infected group at the corresponding time point of infection (Fig. 6L).

## 4. Discussion

As defined by the World Health Organization (WHO), Post-COVID syndrome (PCS) is a condition that encompasses fatigue, shortness of breath, cognitive dysfunction, and other symptoms due to previous SARS-CoV-2 infection that cannot be explained by another diagnosis. Symptoms usually appear 3 months after the onset of COVID-19 and last for at least 2 months (Soriano et al., 2022). Post-COVID syndrome has been reported to affect between 7.5% and 89% of COVID-19 patients (Chen et al., 2022; Nasserie et al., 2021; Subramanian et al., 2022). It is noteworthy that PCS is not limited to severe cases, as 29.6% of patients with mild illness are affected by this condition (Cazé et al., 2023). Considering that most COVID-19 patients (about 80%) develop mild disease (Huang et al., 2020), it is of great importance to study this condition in mild models of coronavirus infection. Here, through a robust characterization of MHV-A59 lung infection and central nervous system dysfunctions, we showed, to the best of our knowledge for the first time, that intranasal instillation of murine betacoronavirus MHV-A59 induced: (I) transient infection and mild lung disease in wild-type C57BL/6J mice; (II) more severe inflammatory response in the lung of female mice, characterized by a robust T cell response; (III) behavioral and cognitive changes, mainly in female mice, that persisted for up to 2 months; (IV) more severe inflammatory response in the brain of female mice (comparing with male counterparts) characterized by neutrophil infiltrate, microglial activation, and IFN-γ response; (V) female hormone-dependent disease phenotype on brain and behavioral sequelae.

The emergence of SARS-CoV-2 and the risk of new coronavirus outbreaks, which had been overlooked for decades (Morens et al., 2004), has emphasized the need to develop models to better understand the pathogen-host interaction and to evaluate potential therapeutic treatments and vaccines for coronavirus infections. Intranasal instillation of different MHV strains in mice has been used as an *in vivo* platform to study coronavirus lung disease (Andrade et al., 2021; Pimenta et al., 2023; Queiroz-Junior et al. 2023; De Albuquerque et al., 2006; Yang et al., 2014). In a prior study, Yang et al. characterized MHV-A59 lung infection in C57BL/6 mice as a model for acute respiratory distress syndrome (Yang et al., 2014). MHV-A59 was able to induce acute lung inflammation in C57BL/6 mice with leukocyte infiltration and hemorrhage, as well as increased mRNA expression of pro-inflammatory mediators such as CXCL10, IFN-γ, TNF, and IL-1β (Yang et al., 2014). Here, MHV-A59 infection also induced mild clinical pulmonary signs, with mild and transient histopathologic lung changes. Mice also exhibited acute lung dysfunction, as evidenced by decreased lung volume and compliance. This stiff behavior of the respiratory system is consistent with pulmonary restrictive disease (Mortola 2019), and COVID-19 patients also show impairment in lung function, including those with mild respiratory symptoms (Mo et al., 2020; Altmann et al., 2023). Fibrosis is an important factor associated with impaired lung function, and COVID-19 patients showed abnormal collagen deposition in lung tissue (Schaller et al., 2020). Despite the increased levels of TGF-β, a key regulator of collagen deposition (Meng et al., 2016), in the lungs of MHV-A59-infected mice at post-acute time points, we did not observe pulmonary fibrosis throughout the course of the disease. Together with the recovery of lung volume and compliance 30 days post-infection, this suggests that the impaired lung function observed in MHV-A59-infected mice is primarily related to the acute phase of lung disease rather than persistent structural changes in lung tissue.

The immune response against MHV-A59 infection had a distinct sex-related phenotype. Indeed, several pieces of evidence suggest that biological sex is an important factor that impacts COVID-19 immune response, outcome, and development of PCS symptoms (Scully et al., 2020; Takahashi et al., 2020). Here, female MHV-A59 infected-mice had a robust lung T cell response, with higher levels of IFN-γ compared to male infected-mice. Conversely, male MHV-A59 infected-mice had prolonged levels of pulmonary chemokines involved in innate immune cell recruitment, such as CCL3, CCL5 and CXCL1. Accordingly, in COVID-19 patients, males have a more robust induction of innate immune response, with higher levels of IL-8, IL-18, and CCL5, and accumulation of non-classical monocytes in lung tissue, while female patients have more abundant activated T cells (Takahashi et al., 2020). This highlights that female and male MHV-A59 infected mice may differ regarding innate and adaptive immune response in the lung tissue, as observed in COVID-19 patients.

Mice infected with MHV-A59 completely cleared the virus in the lung tissue, indicating the host’s ability to cope with the infection. In addition, replicating MHV-A59 could not be detected in the plasma or spleen but in the liver, and also viral RNA copies in the brain, suggesting limited viral spread to extrapulmonary sites. In terms of systemic changes, MHV-A59 infection leads to a transient increase in NLR in the blood, which is also seen in COVID-19 patients and is an indicator of disease severity and mortality (Henry et al., 2020; Liao et al., 2020; Ponti et al., 2020). Of note, mice fully recovered from the imbalance in NLR mediated by MHV-A59 infection. Therefore, this model was characterized by mild lung disease with mild systemic alterations and viral dissemination, constituting a suitable platform to study sequelae post-coronavirus infection. Importantly, infection with MHV-1, another murine betacoronavirus, was recently shown to recapitulate some aspects of PCS in mice (Masciarella et al., 2023).

In the current model, female mice infected with MHV-A59 showed acute olfactory discrimination dysfunction. Olfactory loss is a core symptom of acute COVID-19 (Harapan and Yoo, 2021) as well as PCS (Winter et al., 2023). MHV-A59 infection also triggered motor impairment in female mice, affecting muscular strength of the forelimbs in both sexes at 16-dpi. This feature has recently been reported in patients with PCS (Ramírez-Vélez et al., 2023). In addition, both female and male mice presented anhedonic-like behavior, corroborating previous studies with PCS patients (Lamontagne et al., 2021; Sayed et al., 2021). Interestingly, only female mice demonstrated impairment in spatial working memory and short-term aversive memory in the Y-maze test and in the step-down inhibitory avoidance test, respectively, at 60-dpi. These cognitive deficits have been more commonly observed in women with long-COVID compared to men (Bai et al., 2022). Hence, the present model provides, for the first time, supporting evidence for the predominance of behavioral and cognitive impairments observed in PCS, particularly among females.

Previous findings from our research group demonstrated that synaptosomes from MHV-3 infected mice have increased glutamate release and intracellular calcium levels (Pimenta et al., 2023). In the current model, MHV-A59-infected female mice also had significant glutamate release and increased intracellular Ca^2+^ levels in the hippocampus at 30-dpi. This is suggestive of neuronal excitotoxicity, as viral infections, including HIV, ZIKA and H1N1, have already been shown to impair glutamatergic transmission, thus impairing neural signaling (Costa et al., 2017; Düsedau et al., 2021; Gorska and Eugenin, 2020). Importantly, a greater number of microglia/macrophage (IBA-1) and astrocytes (S100B) were found in the cerebral cortex and hippocampus of female mice when compared to male mice. The increased number of these cells is suggestive of enhanced neuroinflammation and possibly neurodegeneration (Kwon and Koh, 2020; Vandenbark et al., 2021). Moreover, microglia of infected female mice but no male mice showed high expression of inducible nitric oxide synthase (iNOS). Nitric oxide (NO) is essential in synaptic transmission and brain plasticity, mainly in the cortex and hippocampus. However, high levels of NO and nitrergic/oxidative stress can lead to synaptic impairment and early neurodegeneration (Balez and Ooi, 2016).

The more robust neuroinflammation in female mice was further confirmed by detection of immune cells in the brain. MHV-A59-infected female mice presented increased number of IFN-γ^+^-releasing CD4^+^ T lymphocytes in the brain, with Ki67^+^ (cell proliferation marker) and CD69 (cell activation marker) expression, aside higher numbers of Ki67^+^ CD8^+^ T cells in relation to males. T cells are especially important for limiting viral replication, as they can gain access to the brain parenchyma through local recognition of viral antigens by T cell receptors (Steinbach et al., 2016). The interaction between microglia and T cells is also crucial within the CNS parenchyma, since the effector functions of these lymphocytes depend on this communication (Ai and Klein, 2020). However, when this response is not finely regulated, the results can be deleterious. A study demonstrated that T cells might be associated with neurocognitive sequelae in surviving animals during neuropathogenic viral infections, such as ZIKV and West Nile Virus (WNV). This occurs mainly through the signaling of IFN-γ released by specific CD8^+^ T cells infiltrating the CNS, which induces the activation of microglia (Garber et al., 2019). This microglial activation is correlated with several neurotoxic effects, such as excessive complement-mediated synapse elimination, neurodegeneration and decreased adult neurogenesis (Klein et al., 2019). WT mice infected with MHV V5A13.1 presented CNS inflammation and demyelination significantly less severe than T CD4^−/−^ mice (Lane et al., 2000). Regarding other inflammatory mediators, MHV-A59 infection did not alter IL-6 levels in the PFC of female mice compared to controls. In males, IL-6 levels increased at 30- and 60-dpi. However, in the hippocampus, females exhibited increased IL-6 at 8-dpi compared to the mock group and infected males. SARS-CoV-2-induced IL-6 may influence working memory and cognitive systems (Alnefeesi et al., 2021). Increased expression of systemic IL-6 induced by SARS-CoV-2 infection can cross the blood-brain barrier to activate microglia and interfere with memory (Alnefeesi et al., 2021; Vos et al., 2022). Although IL-6 is mostly characterized by its proinflammatory profile, it has been demonstrated that it also participates in neurogenesis, in the response of mature neurons and glial cells in conditions of brain homeostasis (Erta et al., 2012). It is possible that, in the acute infection, especially in females, IL-6 is exerting its pro-inflammatory role in response to MHV-A59, while its later increase in males appears to be a compensatory/protective response to the system to return to homeostasis.

In addition to the inflammatory burden, MHV-A59 triggered an impairment in neurotrophic factors. MHV-A59 infection decreased BDNF levels in the PFC in females at 8- and 16-dpi and in males at 8-dpi. In the hippocampus, females exhibited reduction from 2-to 16-dpi, while males showed this reduction only from 2-to 8-dpi. Such delayed recovery of BDNF in infected females may have contributed to the neuropsychiatric symptoms. BDNF is a key factor in survival, synaptic plasticity, and reorganization of the brain microenvironment (Chen et al., 2020). BDNF may play an antidepressant role in the PFC (Li et al., 2018) and hippocampus (Li et al., 2017), besides reducing anxiety-like behavior in rats (Cirulli et al., 2004). The levels of CX3CL1 showed a general pattern of increase followed by decrease in the PFC and hippocampus of female and male mice, suggesting a homeostatic response to control the acute effects of infection. The CX3CL1/CX3CR1 axis is important for neuron-microglia communication and the predominant function of CX3CL1 in the CNS is believed to reduce the pro-inflammatory response (Subbarayan et al., 2022). Furthermore, impairment of this axis has been associated with the development of neuropsychiatric conditions (Chamera et al., 2020).

Biological sex affects both innate and adaptive immune responses to infections and vaccination (Klein and Flanagan, 2016). Accordingly, biological sex has been considered an important factor influencing COVID-19 immune response, outcomes and sequelae (Scully et al., 2020; Sylvester et al., 2022; Takahashi et al., 2020). In view of the worse phenotype of CNS sequelae in female mice after MHV-A59 infection, we investigated the involvement of female sex-hormones. There is evidence that SARS-CoV-2 spike protein binds and modulates estrogen receptors (Solis et al., 2022). Infected mice presented an increase in estradiol at 60 dpi. Estrogen signaling is associated with sex differences (Dhakal et al., 2021; Kocanova et al., 2010) while the potential protective effects of estrogens on acute COVID-19 have been widely debated in the literature (Al-kuraishy et al., 2021). Most epidemiological studies on COVID-19 have shown a sex-related mortality trend, with men being more vulnerable (Solis et al., 2022). Male laboratory animals are also more susceptible to SARS-CoV and SARS-CoV-2 infection compared to females (Channappanavar et al., 2017; Dhakal et al., 2021; Ruiz-Bedoya et al., 2022).

Regarding PCS, women seem to be more likely to develop post-COVID syndrome (Bai et al., 2022). Female hormones have a protective effect in acute COVID-19, but may play a role in maintaining the hyperinflammatory state of the acute phase, even after recovery (Bienvenu et al., 2020; Mohamed et al., 2021). In a female hormone-deficient environment triggered by ovariectomy, we showed that female mice had similar lung lesions of SHAM-infected mice, but excitotoxicity, behavioral and cognitive impairments were significantly prevented. In line with these results, an experimental study demonstrated that treatment with estradiol did not minimize pulmonary complications in male hamsters infected with SARS-CoV-2 (Dhakal et al., 2021). Our model, in general, mimics several PCS signs and clearly shows a sex-dependent phenotypic susceptibility in agreement with epidemiological data. However, further mechanistic studies are needed to answer how such female hormone deficiency can prevent the CNS sequelae induced by coronavirus infection. Herein, our data suggested that estradiol may play a role in promoting cognitive disturbances. However, it is crucial to broaden the scope of the investigation to include other hormones, such as progesterone, testosterone, and gonadotropins. This comprehensive approach will enable a more thorough understanding of the hormonal influences in the PCS model.

In conclusion, this is the first time that an experimental study recapitulates several aspects of the human Post-COVID syndrome, clearly showing the sex differences regarding cognitive and behavioral outcomes. MHV-A59 induced mild acute lung inflammation and triggered chronic neuropsychiatric and musculoskeletal disorders, which were dependent on female hormones. This model offers a distinct platform to study the pathogenesis of PCS within a BSL2 structure. Its significance lies in addressing the challenge of studying behavioral and cognitive impairment in BSL3 laboratories mandated for SARS-CoV-2. Additionally, it facilitates the exploration of the therapeutic efficacy of antiviral, anti-inflammatory, or neuroprotective strategies, enhancing the overall management of COVID-19.

## Funding

This study was funded by the Fundação de Amparo à Pesquisa do Estado de Minas Gerais grant numbers: APQ02281-18, APQ-00826-21; APQ-01078-21, APQ-02402-23 and APQ02618-23. Conselho Nacional de Desenvolvimento Científico e Tecnológico bythefollowinggrants442731/2020-5; 408482/2022-2, 305932/2022-5 and 422002/2023-2. This work was also supported by grants from Coordenação de Aperfeiçoamento de Pessoal de Nível Superior –CAPES/Brazil (Projeto: CAPES - Programa: 9951 - Programa Estratégico Emergencial de Prevenção e Combate a Surtos, Endemias, Epidemias e Pandemias AUX 0641/2020 - Processo 88881.507175/2020-01) and CAPES 11/2020 Epidemias, N° 88887.506690/2020-00. ThisworkalsoreceivedsupportfromtheNationalInstituteof Science and Technology in Dengue and Host-MicroorganismInteraction (INCT em Dengue), sponsoredbythe Conselho Nacional de Desenvolvimento Científico e Tecnológico (CNPq; Brazil) (Processo CNPQ: 465425 /2014-3) andthe Fundação de Amparo à Pesquisa do Estado de Minas Gerais (FAPEMIG; Brazil) (Processo FAPEMIG: 25036). Andby FINEP - Financiadora de Estudos e Projetos underMCTI/FINEP – MS/SCTIE/DGITIS/CGITS (6205283B-BB28-4F9C-AA65-808FE4450542) grant.

## Declaration of Competing Interest

The authors declare that they have no known competing financial interests or personal relationships that could have appeared to influence the work reported in this paper.

## Acknowledgements

We are grateful to Ilma Marçal de Souza, Rosemeire Oliveira, Letícia Soldati and Tânia Colina for their technical assistance. We also thank the support of the Technological Center for Advanced and Innovative Therapies - CT - Terapias UFMG and the Experimental Platforms of UFMG (flow cytometry - LaboratórioInstitucional de Pesquisa emBiomarcadores (LINBIO) at Pharmacy Faculty/UFMG, Neurological and behavioral analysis - Laboratório de Neurobiologia “Profa Conceição Machado” at ICB/UFMG, *in vivo* platforms of viral infections - GPAV at ICB/UFMG) for providing the facilities and expertise for the conduction of the experiments. VVC and ASM also thanks “Para mulheres na Ciência Prize” provided by L’oréal, UNESCO and ABC. Thanks are also due to the animal biosafety level 3 laboratory at UFMG (Laboratório Institucional de Pesquisa, LIPq); Centro de Laboratórios Multiusuários, CELAM, Laboratório de Biossegurança Nível 3, NB3-ICB.

## Data availability

The data that support the findings of this study are included within the article and are available upon request.

---

**Supplementary figure 1. Intranasal instillation of MHV-A59 in mice results in transitory and mild liver injury.(A)** Histopathological assessment in relation to overall liver inflammatory score. Significance was determined by Two-way ANOVA and Dunnet’s multiple comparison test *p < 0.01 (n = 5-6). **(B)** Representative images of H&E-stained liver sections of mock controls and 5-dpi MHV-A59-infected mice.

**Supplementary** Figure 2. Flow cytometry gates for leukocytes (CD45high), dendritic cells (CD45highCD11b^+^CD11c^+^), T CD4^+^ lymphocytes (CD45highCD3^+^CD4^+^), T CD8^+^ lymphocytes (CD45highCD3^+^CD8^+^), NK cells (CD45highCD3^-^NK^+^), and markers (IFN-γ^+^, Ki67, IL-10)

**Supplementary** Figure 3**. (A)** Hippocampal intrasynaptosomal calcium concentration (n = 5-7) and **(B)** glutamate release. Significance was determined by Student’s t-test to data that passed Shapiro-Wilk test and Mann–Whitney test to data that did not pass Shapiro-Wilk test *p < 0.05(n = 6-8). **(C)** Brain histopathological score. Significance was determined by Two-way ANOVA and Dunnet’s multiple comparison test to analyze the differences between mock controls and infected mice of respective sex at different time points. Significance was determined by Two-way ANOVA and the Šídák multiple comparison test to analyze the differences between the sexes at the respective infection time points. *p < 0.01 (n = 3-8). **(D)** Representative images of H&E-stained brain sections of mock controls and 8-dpi MHV-A59-infected mice.

**Supplementary** Figure 4. Flow cytometry gates for leukocytes (CD45high), activated microglia (CD45intCD11b^+^F4/80^+^MHCII^+^), T CD4^+^ lymphocytes (CD45highCD3^+^CD4^+^), T CD8^+^ lymphocytes (CD45highCD3^+^CD8^+^), and markers (IFN-γ^+^, CD69, Ki67).

**Supplementary** Figure 5**. (A)** Representative images of H&E-stained lung sections of sham and OVX mock controls and 5-dpi MHV-A59-infected mice. **(B)** Representative images of H&E-stained brain sections of sham and OVX mock controls and 5-dpi MHV-A59-infected mice.

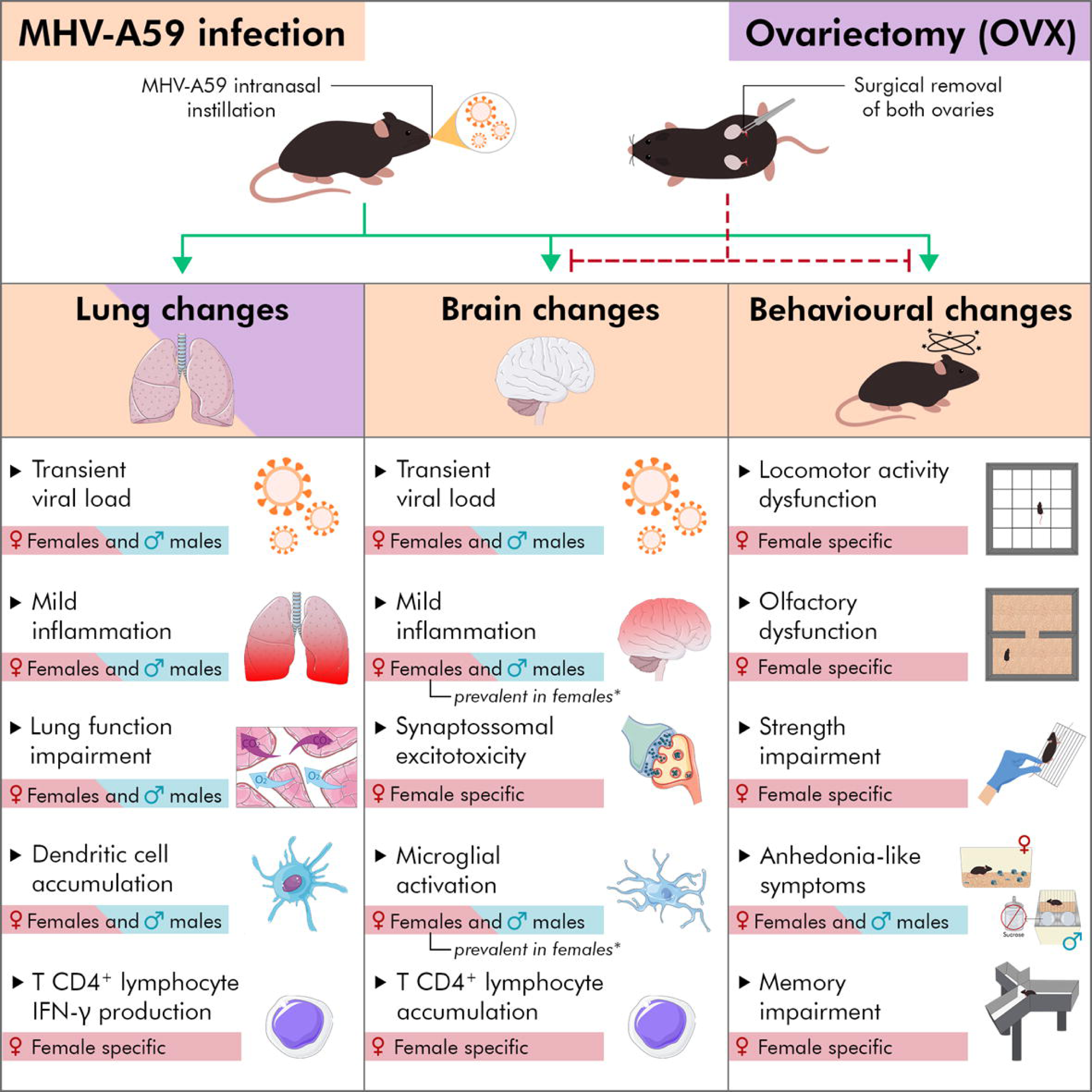

## Notes

### Competing Interest Statement

The authors have declared no competing interest.

