## Supplementary figures and images for "Neuropsychiatric sequelae in an experimental model of post-COVID syndrome in mice"

### Figure S1

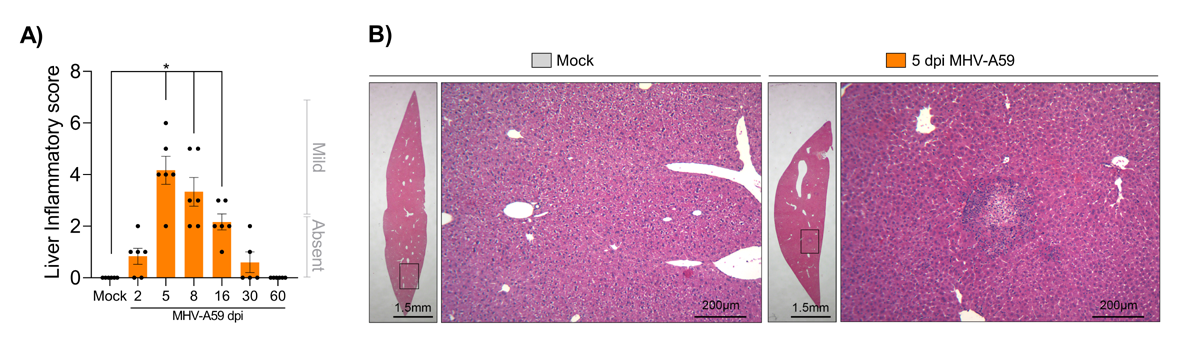

### Figure S2

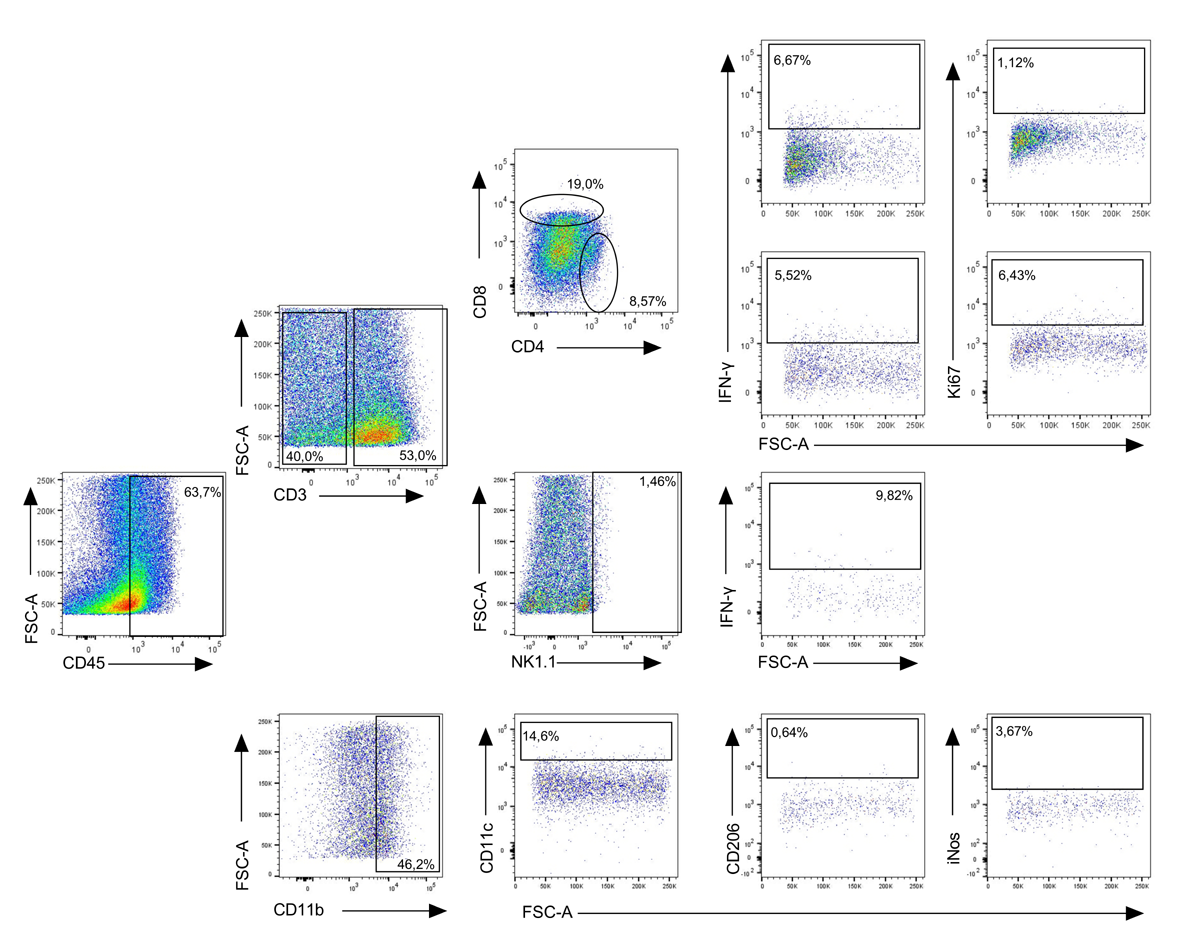

### Figure S3

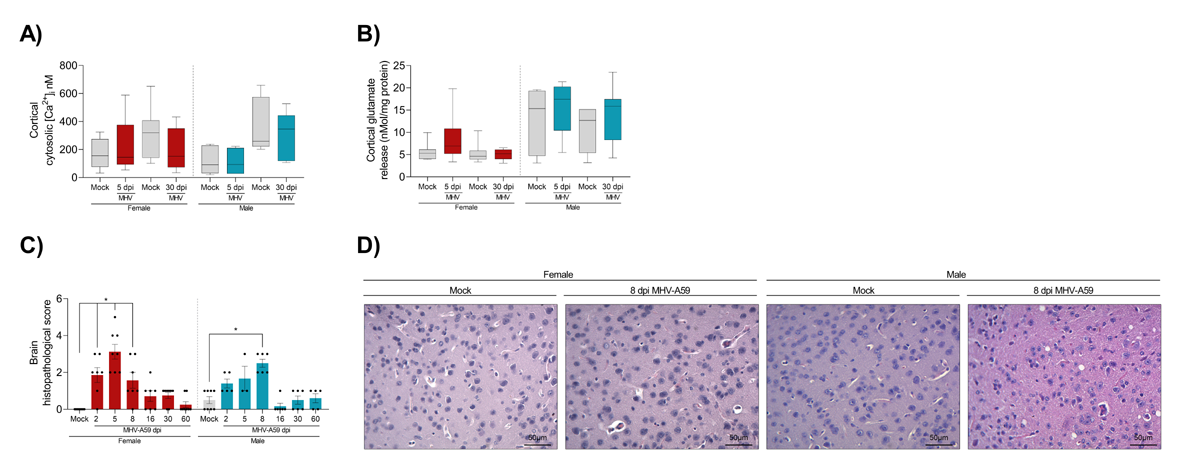

### Figure S4

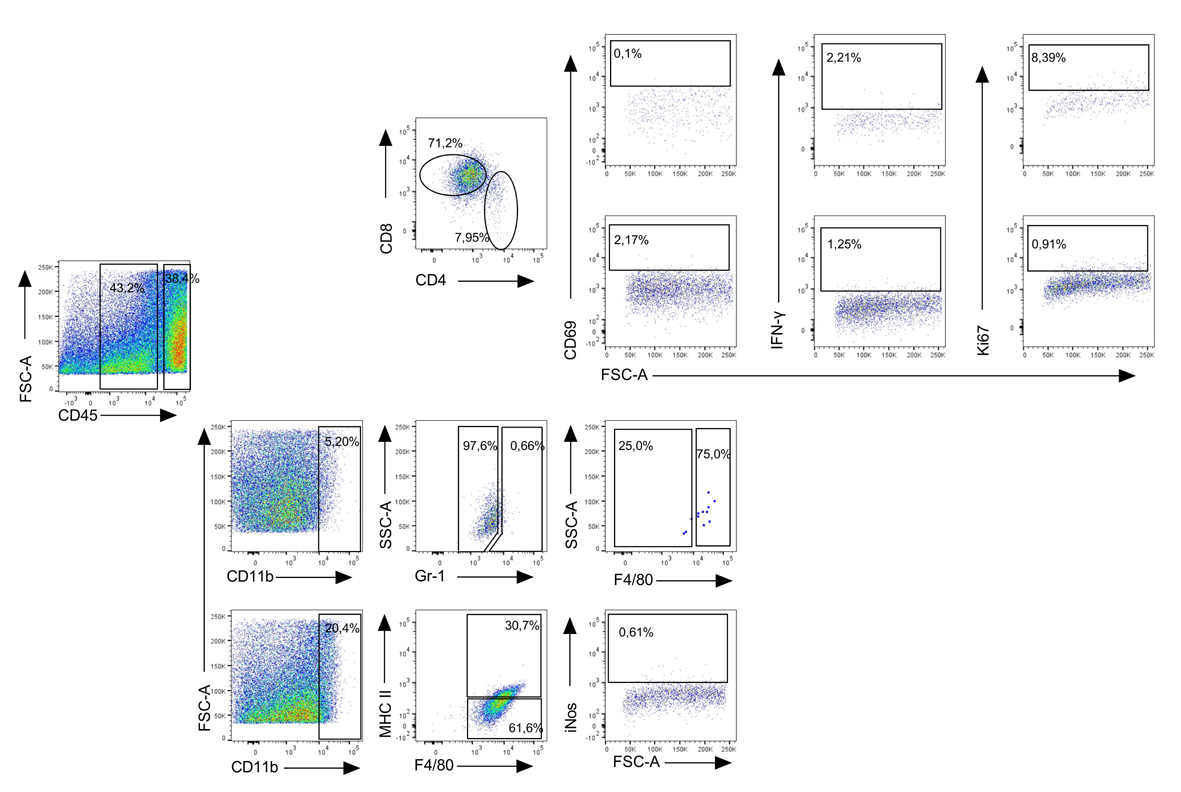

### Figure S5

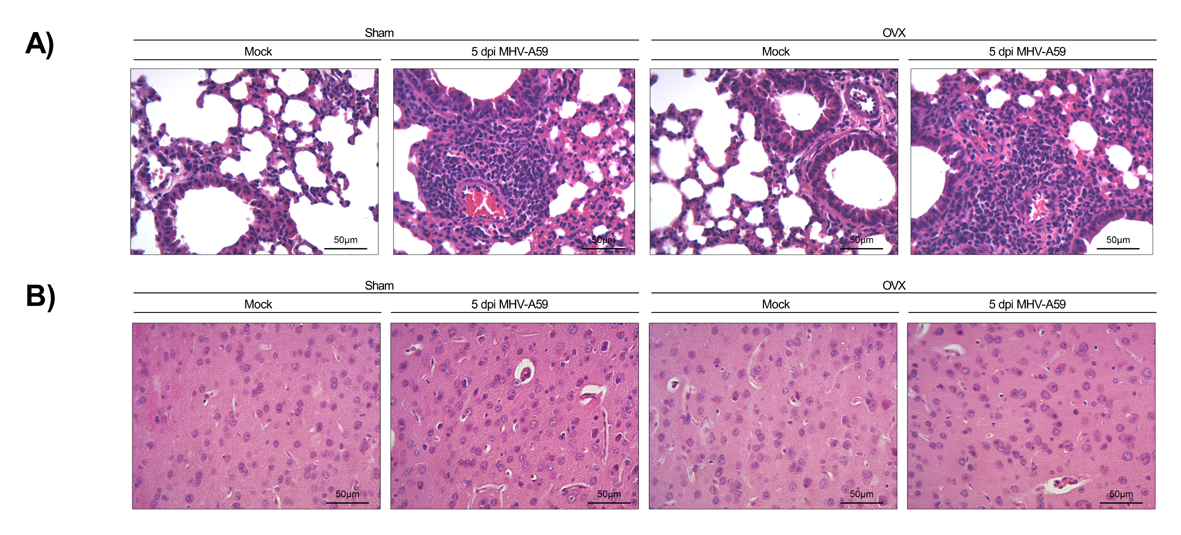
